# Cryo-EM structure of the Inner Ring from *Xenopus laevis* Nuclear Pore Complex

**DOI:** 10.1101/2021.11.13.468242

**Authors:** Gaoxingyu Huang, Xiechao Zhan, Chao Zeng, Ke Liang, Xuechen Zhu, Yanyu Zhao, Pan Wang, Qifan Wang, Qiang Zhou, Qinghua Tao, Minhao Liu, Jianlin Lei, Chuangye Yan, Yigong Shi

## Abstract

Nuclear pore complex (NPC) mediates nucleocytoplasmic shuttling. Here we present single-particle cryo-electron microscopy (cryo-EM) structure of the inner ring (IR) subunit from *Xenopus laevis* NPC at an average resolution of 4.4 Å. The symmetric IR subunit comprises a cytoplasmic half and a nuclear half. A homo-dimer of Nup205 resides at the center of the IR subunit, flanked by two molecules of Nup188. Four molecules of Nup93 each places an extended helix into the axial groove of Nup205 or Nup188, together constituting the central scaffold. The channel nucleoporin heterotrimer (CNT) of Nup54/58/62 is anchored on the central scaffold. Six Nup155 molecules interact with the central scaffold and together with the NDC1-ALADIN hetero-dimers anchor the IR subunit to the nuclear envelope and to outer rings. The scarce inter-subunit contacts may allow sufficient latitude in conformation and diameter of the IR. Our structure of vertebrate IR reveals key insights that are functionally important.

---

NPC is present on the nuclear envelope (NE) of all eukaryotic cells and mediates nucleocytoplasmic transport (*1, 2*). A vertebrate NPC consists of four circular scaffolds: a cytoplasmic ring (CR), an inner ring (IR), a nuclear ring (NR), and a luminal ring (LR) (*3-5*). IR resides in the central layer of the NPC and bridges the outer rings CR and NR. Of the four rings, only the LR resides in the lumen of the NE. The cytoplasmic filaments (CF) are connected to the CR and facilitate nucleocytoplasmic transport. Structural investigation of the NPC has yielded a wealth of information on individual nucleoporins, subcomplexes, and ring scaffolds of the NPC (*2, 4, 6, 7*).

A vertebrate NPC has an extraordinary molecular mass and displays considerable conformational flexibility despite its 8-fold symmetry. These aspects have made cryo-EM analysis of the NPC through single particle analysis (SPA) technically challenging. Consequently, cryo-electron tomography (cryo-ET) through sub-tomogram averaging (STA) has been the dominant approach in the structural investigation of NPC during the past two decades. The best resolution achieved for cryo-ET reconstruction of the IR subunit is approximately 21 Å for the human NPC (*8*). This EM map allowed generation of composite coordinates for human NPC through docking of known X-ray structures (*8*).

We took the cryo-EM SPA approach and succeeded in the reconstruction of the CR subunit from *Xenopus laevis (X. laevis)* NPC with a local resolution of about 5-8 Å (*9*). More recently, relying on enhanced methods of sample preparation and cryo-EM analysis, we have markedly improved the local resolution of the *X. laevis* CR subunit to 3.8 Å (*10*). Detailed features of secondary structural elements and bulky amino acid side chains are identifiable in a portion of the core region.

In this manuscript, we report the cryo-EM reconstruction of the IR subunit of the *X. laevis* NPC at an average resolution of 4.4 Å, with local resolution reaching 4.0 Å. The EM maps allow docking of known and predicted structures of *X. laevis* nucleoporins (*10, 11*) as well as identification of secondary structural elements. Based on the EM maps, we have generated atomic coordinates for 30 nucleoporins of the *X. laevis* IR subunit, which include 19,315 amino acids. This structure reveals the underpinnings of vertebrate IR subunit assembly.

## Cryo-EM analysis of the *X. laevis* IR subunit

The same cryo-EM datasets used for reconstruction of the CR subunit from *X. laevis* oocytes (*10*) were used for reconstruction of the IR subunit. A total of 800,825 NPC particles were manually selected from 33,747 micrographs (Fig. S1*A)*. With C8 symmetry, the intact IR was reconstructed at an initial resolution of 22 Å using 660,302 NPC particles (Fig. 1*A*; Fig. S1*A,B)*, allowing extraction of the IR subunits (Fig. S2*A)*. Using data at the bin-2 level, we generated a reconstruction of the IR subunit at 5.6 Å resolution. The refined subunits were projected back to the original IR, followed by subunit re-extraction (Fig. S2*B)*. Using data at the bin-1 level, 2,093,631 particles yielded the final reconstruction for the IR subunit at an average resolution of 4.4 Å (Fig. 1*B,C*; Fig. S3-S7; Tables S1 & S2). The local resolution reaches 4.0 Å.

**Fig. 1.**
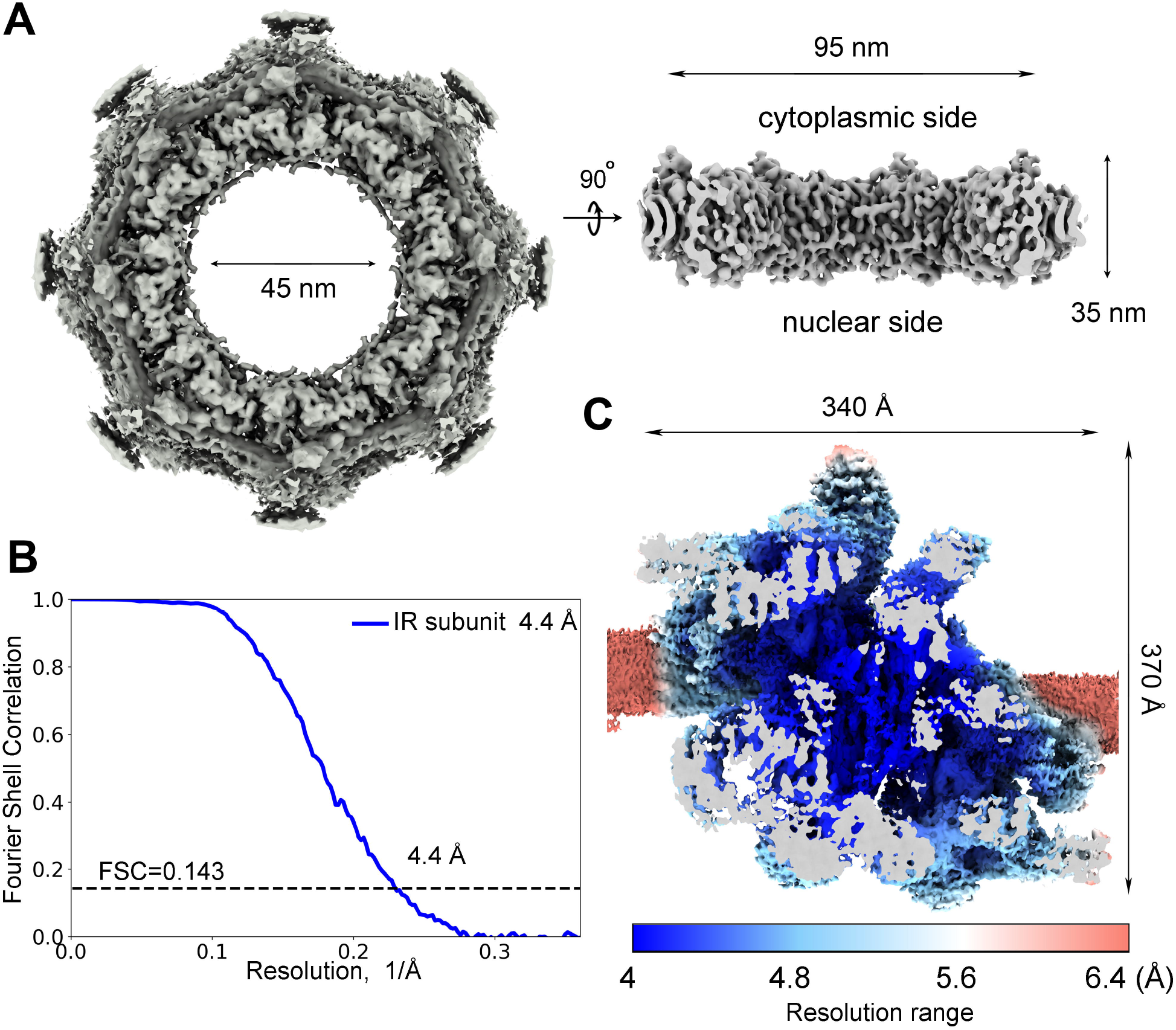
Cryo-EM structure of the inner ring (IR) of the *X. laevis* NPC. (*A*) Cryo-EM reconstruction of the IR of *X. laevis* NPC at 22 Å resolution. Two perpendicular views are shown. The inner diameter of the IR is approximately 45 nm. (*B*) The final EM reconstruction for the IR subunit displays an average resolution of 4.4 Å. Shown here is a curve of the Fourier Shell Correlation (FSC) over resolution. (*C*) Distribution of the local resolution for the EM reconstruction of the IR subunit. The color-coded resolution range is shown below.

Compared to previous studies, our cryo-EM reconstruction of the *X. laevis* IR subunit at 4.4-Å resolution constitutes a major improvement. The improved EM density maps allowed unambiguous assignment of most components in the IR subunit and accurate placement of secondary structural elements. We identified 30 molecules of nine distinct nucleoporins in each IR subunit, including four copies of Nup93, six copies of Nup155, four copies of CNT (each comprising Nup62/Nup58/Nup54) (*12*), and two copies each for Nup205, Nup188, NDC1, and ALADIN.

The atomic coordinates of Nup93 and Nup205 from the CR subunit (*10*) were docked into the EM map with little adjustment. Using AlphaFold (*11*), we generated atomic coordinates for each of the other *X. laevis* nucleoporins. We individually docked each predicted structure into the 4.4-Å EM map and made minor adjustment (Fig. S4-S7). Compared to the core region, the EM density in the peripheral region of the IR subunit has a lower resolution (Fig. S7). Nonetheless, the AlphaFold-predicted structure of the ALADIN-NDC1 hetero-dimer fits the EM density reasonably well. The final atomic model of the *X. laevis* IR subunit contains 19,325 amino acids, with 788 α-helices and 312 β-strands (Fig. S8-S13; Table S3).

### Overall structure of the IR subunit

The IR subunit displays a two-fold symmetry, with a cytoplasmic half and a nuclear half separated along the NE (Fig. 2*A*). At the center of the IR subunit, two molecules of Nup205 form a homo-dimer, which is flanked by two Nup188 molecules (Fig. 2*B*; Fig. S14). Four molecules of Nup93 closely interact with the Nup205/Nup188 hetero-tetramer, with each Nup93 placing an extended helix into the axial groove of Nup205 or Nup188 (Fig. S14). Together, eight molecules of Nup93, Nup188, and Nup205 assemble into a central scaffold of the CR subunit.

**Fig. 2.**
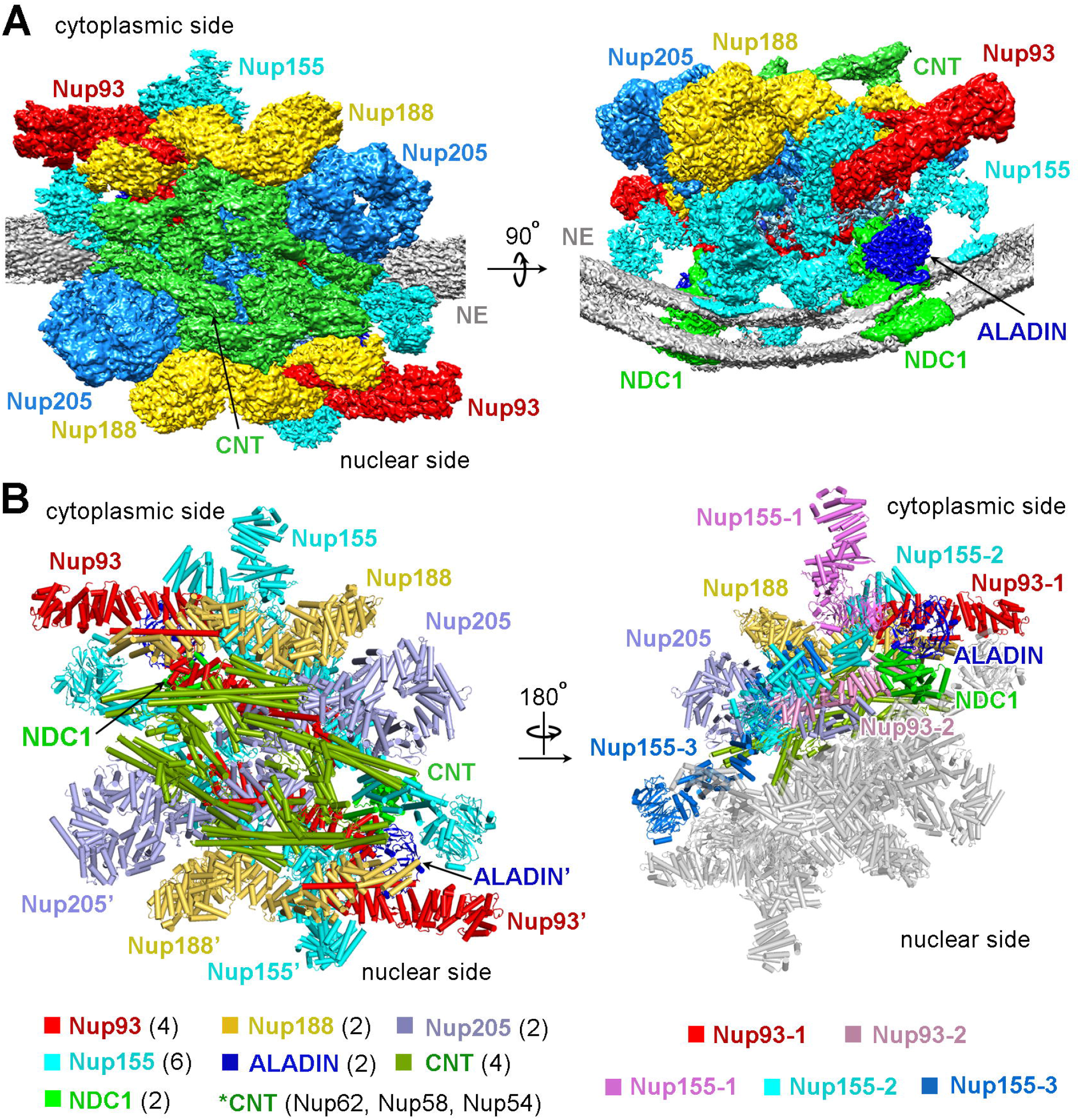
Overall structure of the IR subunit from the *X. laevis* NPC. (*A*) Overall EM density map of the IR subunit from *X. laevis* NPC. Two perpendicular views are shown. The IR subunit displays a 2-fold symmetry, with a cytoplasmic half and a nuclear half. The EM maps for individual nucleoporins in each half are color-coded identically. The NE is shown in grey. (*B*) Structure of the IR subunit from *X. laevis* NPC. Two views are shown. In the left panel, individual nucleoporins are color-coded. In the right panel, nucleoporins of the nuclear half are colored gray; different copies of the same protein in the cytoplasmic half are differentially colored.

The central scaffold associates with four CNTs on the side of the central pore of the NPC (Fig. S14). This is accomplished in part by an N-terminal helix of Nup93, which interacts with triple helical bundles of Nup62, Nup58, and Nup54 (*12*). The central scaffold also associates with six molecules of Nup155 and two hetero-dimers of ALADIN-NDC1 on the side of the NE. Together, 10 molecules of Nup155, ALADIN, and NDC1 assemble into a substructure that has a concave curvature to contact the convex surface of the NE (Fig. S14). Notably, all six β-propeller domains of Nup155 and two ALADIN β-propellers are located in close proximity to the NE. NDC1 contains a pore domain (PD) and a transmembrane domain (TM). The PD interacts with ALADIN (*13, 14*); the TM helps anchor the IR on the NE (Fig. S14).

For ease of discussion, three Nup155 molecules on the cytoplasmic side are designated Nup155-1, Nup155-2, and Nup155-3 along the cytonuclear axis of the IR subunit (Fig. 2*B*; Fig. S14). Similarly, two Nup93 molecules on the cytoplasmic side are referred to as Nup93-1 and Nup93-2, with the latter closer to the center of the IR subunit. Each of the corresponding nucleoporins on the nuclear side is named with an apostrophe. The two halves of the IR subunit are nearly identical to each other, with a root-mean-squared deviation (RMSD) of about 1.34 Å over 8,911 Cα atoms (Fig. S15). Structural comparison of individual nucleoporins between the cytoplasmic and nuclear halves reveals few differences (Figs. S16 & S17). We therefore choose to discuss packing interactions mainly in the cytoplasmic half of the IR subunit.

### Nup188 and Nup205 as the central components

It was unclear whether Nup188 and Nup205 are both present in the IR. Based on the EM map, we conclusively identified two molecules of Nup188 and two molecules of Nup205 in the IR subunit (Fig. 3*A*; Fig. S4). Nup188 and Nup205 are similar in both size and overall fold (Fig. 3*B*). At the center of the IR subunit, Nup205 and Nup205’ form a symmetric homo-dimer that has the appearance of a dumbbell, with their C-terminal helices facing each other at the center (Fig. 3*A*). Specifically, two helices α75/α78 at the C-terminus of Nup205 stack against two corresponding helices of Nup205’ at a roughly perpendicular angle (Fig. 3*C*). Nup205 directly associates with Nup188. Specifically, the surface loop between helices α55 and α56 of Nup205 is positioned in close proximity to helix α33 of Nup188 (Fig. 3*D*). In addition, the surface loop between α39 and α40 of Nup188 is located close to the surface loop between helices α59 and α60 of Nup205.

**Fig. 3.**
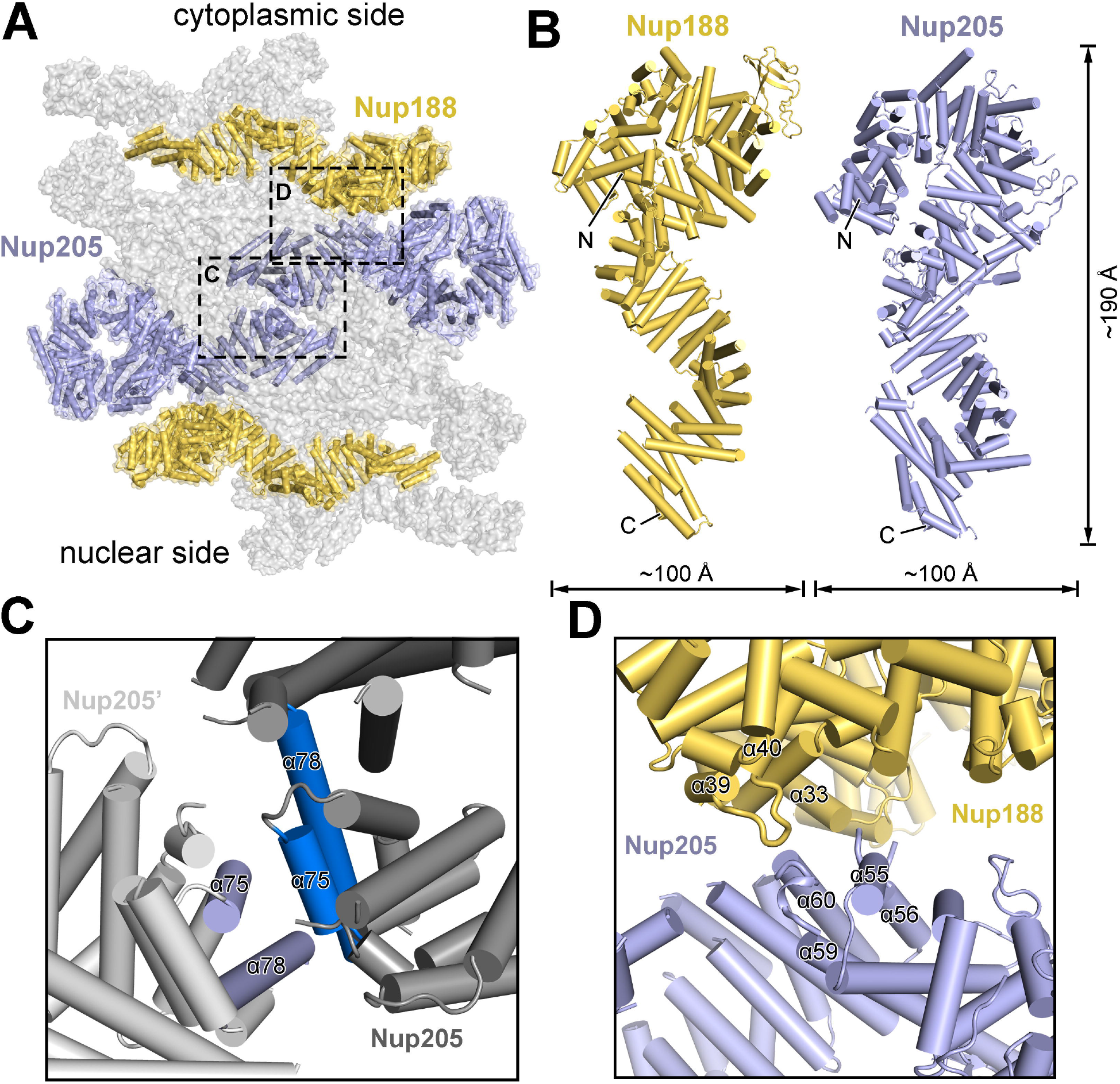
Two molecules of Nup205 and two molecules of Nup188 constitute the central components of the IR subunit. (*A*) A homo-dimer of Nup205 at the center of the IR subunit is flanked by two molecules of Nup188. The two Nup205 molecules contact each other via their C-terminal helices. (*B*) A side-by-side comparison of the structures of Nup188 and Nup205. The overall appearance and size are similar between Nup188 (left panel) and Nup205 (right panel). Nup205 contains a unique TAIL-C domain. (*C*) A close-up view on the dimeric interface between Nup205 and Nup205’. The C-terminal helices α75/α78 from Nup205 stack against the corresponding helices from Nup205’. (*D*) A close-up view on the interface between Nup205 and Nup188. Three inter-helical surface loops from Nup188 and Nup205 are involved in the interactions.

### Formation of the Nup93-Nup188-Nup205 central scaffold

Using the atomic coordinates of Nup93 from the CR subunit (*10*), four molecules were readily placed into the EM density map (Fig. 4*A*; Fig. S5). Notably, Nup93 contains a short α-helix at the N-terminus and an extended N-terminal helix α5 (Fig. 4*B*), which are important for binding CNT and Nup188/Nup205 (*12, 15*), respectively. Four Nup93 molecules form two asymmetric pairs, one each on the cytoplasmic and nuclear sides. Although there is no interaction between the two pairs, two Nup93 molecules within the same pair interact with each other. Specifically, the surface loop between helices α30 and α31 of Nup93-1 may contact helix α14 of Nup93-2 (Fig. 4*C*). In addition, the surface loop between helices α14 and α15 of Nup93-2 is wedged into the crevice formed by helices α35 and α37 of Nup93-1.

**Fig. 4.**
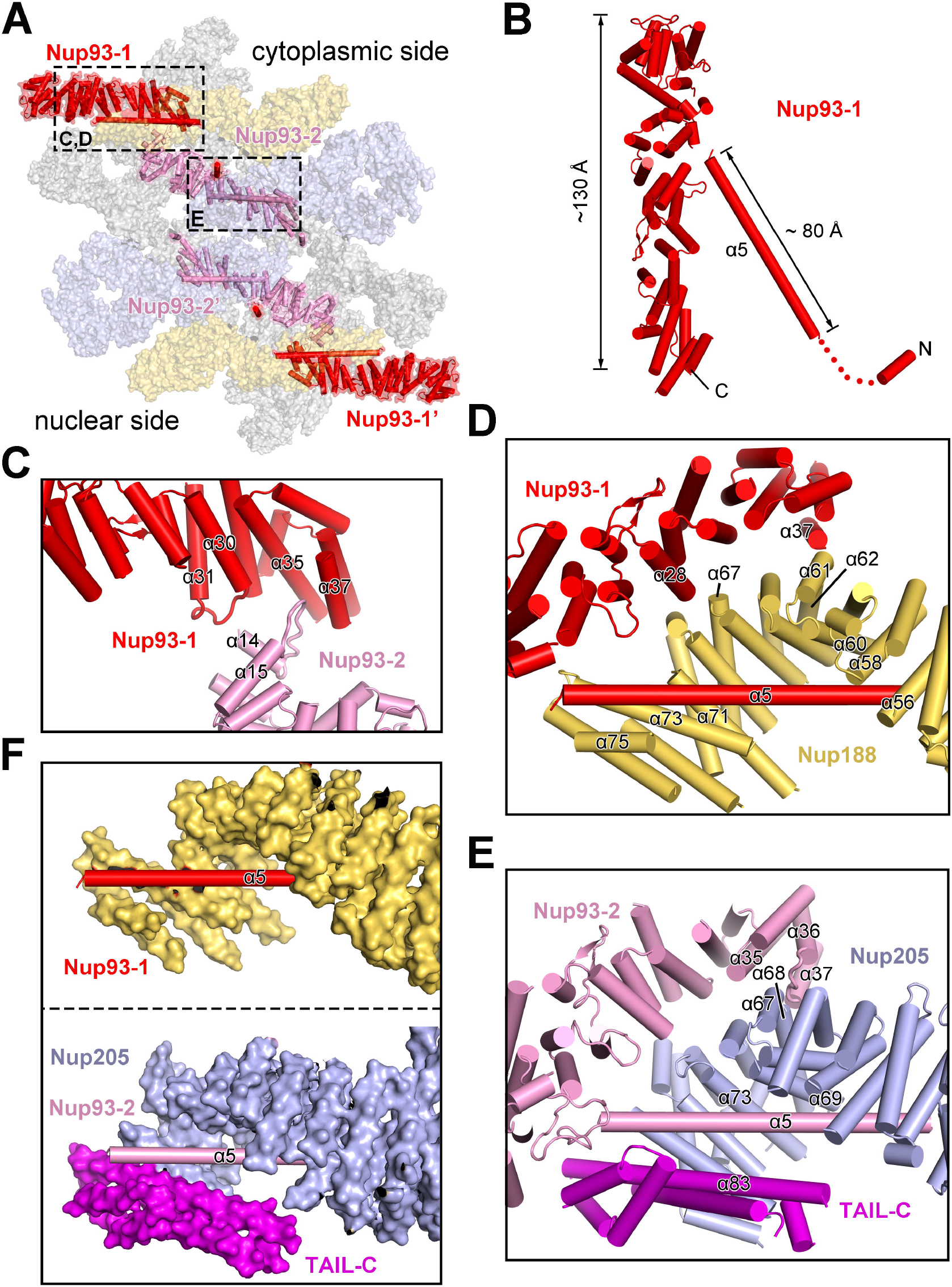
Four molecules of Nup93 interact with Nup188 and Nup205 to form the central scaffold. (*A*) Overall distribution of four Nup93 molecules in the IR subunit. On the cytoplasmic side, Nup93-1 and Nup93-2 interact with Nup188 and Nup205, respectively. (*B*) Structure of Nup93. Shown here is a cartoon representation of Nup93-1. The short N-terminal helix is known to interact with CNT (*12*). The extended N-terminal helix α5 binds Nup188. (*C*) A close-up view on the asymmetric interface between Nup93-1 and Nup93-2. (*D*) A close-up view on the interface between Nup93-1 and Nup188. Notably, helix α5 of Nup93-1 is placed in the axial groove of the Nup188 α-solenoid and interacts with the C-terminal helices of Nup188. (*E*) A close-up view on the interface between Nup93-2 and Nup205. In striking analogy to the Nup93-1/Nup188 interface, helix α5 of Nup93-2 is placed in the axial groove of the Nup205 α-solenoid and interacts with the C-terminal helices of Nup205. (*F*) A side-by-side comparison of the interaction between helix α5 of Nup93 and Nup188 (upper panel) or Nup205 (lower panel). Nup188 and Nup205 are shown in surface representation.

Our EM map reveals previously unknown structural features in the IR subunit. Remarkably, each of the four Nup93 molecules contains an extended α-helix of 80-Å in length at its N-terminus (Fig. S5); this helix α5 traverses the axial groove of the C-terminal portion of the Nup188 or Nup205 α-solenoid (Fig. 4*D*-*F*). Specifically, the N-terminal half of helix α5 from Nup93-1 directly contacts helices α56/α58/α60 from Nup188; the C-terminal half of α5 associates with helices α71/α73/α75 from Nup188 (Fig. 4*D*). In addition, helix α37 of Nup93-1 may contact the surface loop between α61 and α62 of Nup188; helix α28 of Nup93-1 is located close to helix α67 of Nup188.

The interface between Nup93-2 and Nup205 resembles that between Nup93-1 and Nup188. The extended helix α5 from Nup93-2 directly contacts helices α69/α73/α83 from Nup205 (Fig. 4*E*). In particular, helix α83 comes from the TAIL-C domain (*9*), which nearly forms a closed channel with the Nup205 α-solenoid (Fig. 4*F*). In contrast, Nup188 lacks the corresponding TAIL-C and the bound helix α5 from Nup93-1 is more surface-exposed (Fig. 4*F*). In addition, helix α35 of Nup93-2 may contact helix α67 of Nup205 and the surface loop between α36 and α37 of Nup93-2 associates with helices α67/α68 of Nup205 (Fig. 4*E*).

In summary, these four Nup93 molecules form a network of interfaces with the Nup188-Nup205 hetero-tetramer. Together, these eight nucleoporins constitute a central scaffold of the IR subunit, onto which the CNT and the Nup155-ALADIN-NDC1 are anchored.

### Nup155 links the central scaffold

Each Nup155 protein contains an N-terminal β-propeller, followed by an elongated α-helical domain. All six Nup155 molecules in the IR subunit are located on the NE side, with their β-propellers directly contacting the NE (Fig. 5*A*; Fig. S14). Of the six molecules, Nup155-1 and Nup155-1’ directly interact with nucleoporins in the CR and NR, respectively. Specifically, the C-terminal helices of Nup155-1 are sandwiched by inner Nup160 and inner Nup205 from the CR subunit (Fig. 5*B*). These interactions may play a major role in the connection between the IR and the CR/NR.

**Fig. 5.**
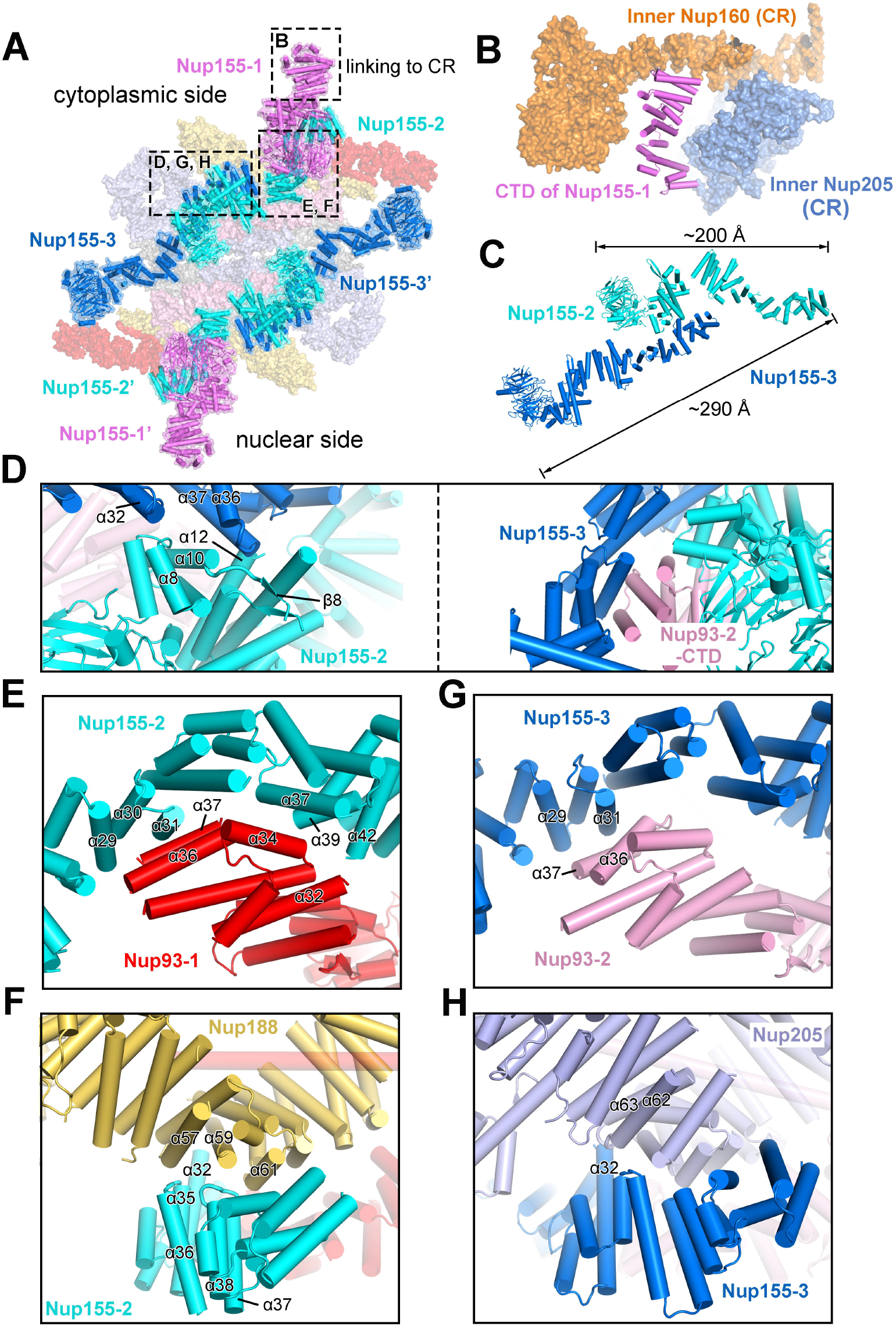
Nup155 links the central scaffold. (*A*) An overall view of six Nup155 molecules in each IR subunit. Of the six Nup155 molecules, Nup155-2 and Nup155-3 form a dimer on the cytoplasmic side; Nup155-2’ and Nup155-3’ form a homo-dimer on the nuclear side. (*B*) Nup155 connects the IR subunit to the CR subunit. The C-terminal helices of Nup155 on the cytoplasmic side of the IR subunit are sandwiched by inner Nup160 and inner Nup205 of the CR subunit. (*C*) A close-up view on the Nup155 homo-dimer. The helical domains of Nup155-2 and Nup155-3 interact with each other in a head-to-tail fashion, generating an elongated Nup155 homo-dimer of about 290 Å in length. (*D*) Close-up views on the interface between Nup155-2 and Nup155-3. Two view are shown. (*E*) A close-up view on the interface between Nup155-2 and Nup93-1. (*F*) A close-up view on the interface between Nup155-2 and Nup188. (*G*) A close-up view on the interface between Nup155-3 and Nup93-2. This interface is analogous to that between Nup155-2 and Nup93-1. (*H*) A close-up view on the interface between Nup155-3 and Nup205. This interface is analogous to that between Nup155-2 and Nup188.

The other four Nup155 molecules form two asymmetric homo-dimers. The elongated α-helical domains of Nup155-2 and Nup155-3 interact with each other in a roughly head-to-tail fashion, creating a homo-dimer of about 290 Å in length (Fig. 5*C*). Helices α8 and α12 of Nup155-2 directly contact helices α32 and α37 of Nup155-3, respectively; the linker sequence between helix α10 and strand β8 of Nup155-2 is in close proximity with the surface loop between α36 and α37 of Nup155-3 (Fig. 5*D*, left panel). Notably, the C-terminal domain (CTD) of Nup93-2 associates with both Nup155-2 and Nup155-3, likely stabilizing their interface (*16*) (Fig. 5*D*, right panel).

This asymmetric dimer interacts with all three components of the central scaffold. The C-terminal helices of Nup155-2 closely stack against the C-terminal helices of Nup93-1 and the middle portion of Nup188. On one hand, helices α29/30/31, α37/α39, and α42 of Nup155-2 may contact helices α36/α37, α34, and α32 of Nup93-1, respectively (Fig. 5*E*). On the other hand, the loop between helices α35/α36, helix α32, and the loop between α37/α38 of Nup155-2 interact with helices α57, α59, and α61 of Nup188, respectively (Fig. 5*F*).

The interactions involving Nup155-3 closely resemble those involving Nup155-2. Nup155-3 closely stacks against Nup93-2 and Nup205. On one hand, helices α29/α31 of Nup155-3 interact with helices α36/α37 of Nup93-2 (Fig. 5*G*). On the other hand, helix α32 of Nup155-3 may directly contact the surface loop between helices α62 and α63 (Fig. 5*H*).

### The ALADIN-NDC1 dimer

The nucleoporin ALADIN, heavily targeted for mutations in triple A syndrome (AAAS) (*17-19*), is thought to directly interact with NDC1 in human cells (*13, 14*). However, the exact locations of ALADIN and NDC1 in the IR subunit have remained enigmatic. In two regions of the EM map next to Nup155-1 and Nup155-1’, there are two lobes of significant density (Fig. 6*A*: Fig. S7). Unfortunately, these regions are in the periphery of the IR subunit where the quality of the EM density is insufficient for *ab initial* identification of nucleoporins. Using AlphaFold (*11*), we generated a predicted structure of the *X. laevis* NDC1; the PD of NDC1 fits the EM map quite well (Fig. S7). Next, we generated an AlphaFold-predicted structure of the ALADIN-NDC1 hetero-dimer, which, to our pleasant surprise, can be docked into the EM density map with little adjustment (Fig. S7).

**Fig. 6.**
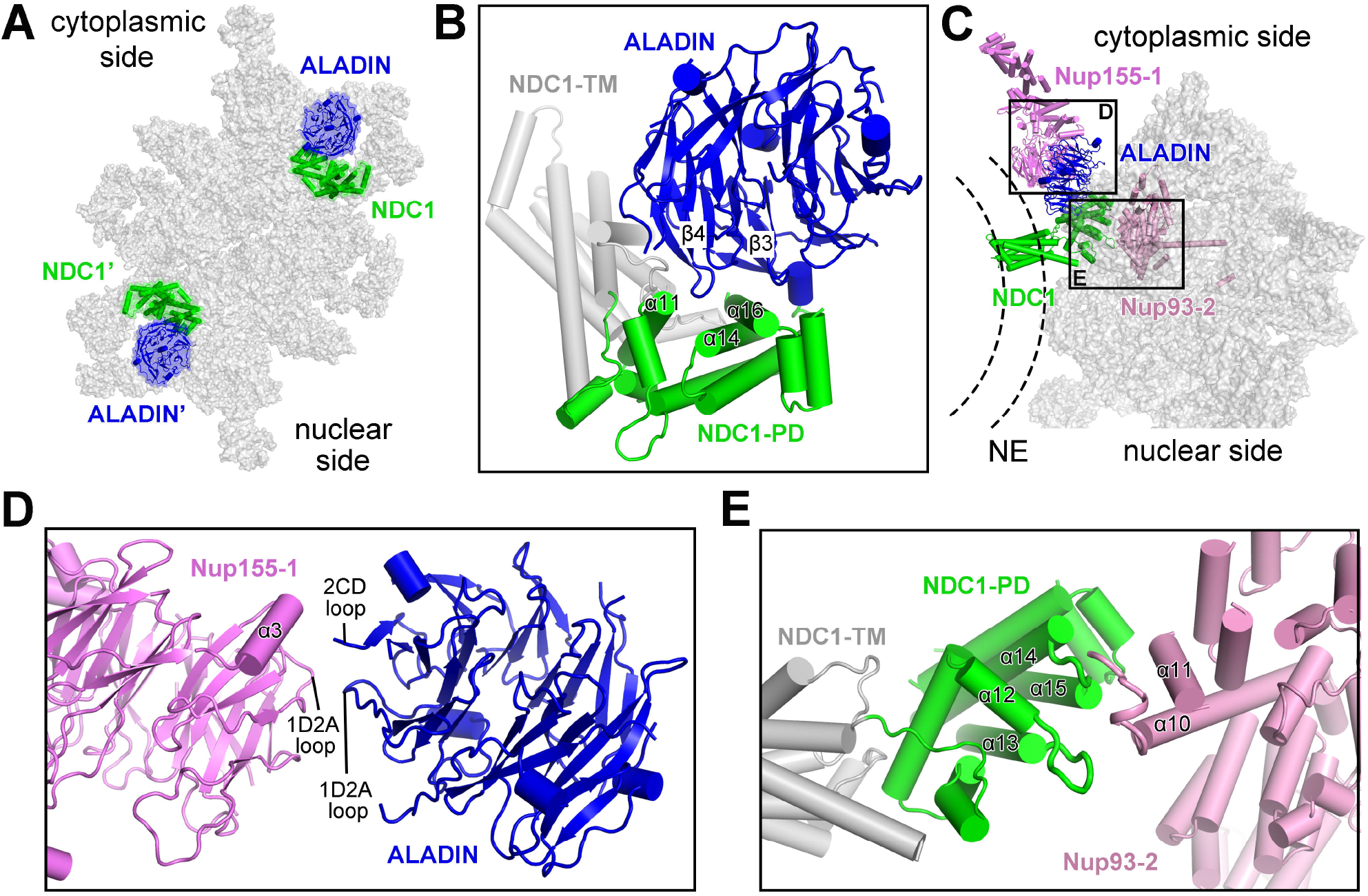
The ALADIN-NDC1 hetero-dimer helps to anchor the IR subunit to the NE. (*A*) An overall view of two ALADIN-NDC1 hetero-dimers in the IR subunit. (*B*) A close-up view on the ALADIN-NDC1 hetero-dimer. Blade-3 and blade-4 of the ALADIN β-propeller are in close proximity to helices α11/α14/α16 of the PD of NDC1. (*C*) An overall view on the cytoplasmic ALADIN-NDC1 hetero-dimer relative to its neighboring nucleoporins Nup155 and Nup93. (*D*) A close-up view on the interface between ALADIN and Nup155-1. (*E*) A close-up view on the interface between NDC1 and Nup93-2.

There are two ALADIN-NDC1 hetero-dimers in each IR subunit, one on the cytoplasmic side and the other on the nuclear side (Fig. 6*A*). In the modeled structure, the loop between strands C and D of blade 4 (known as the 4CD loop) from the ALADIN β-propeller is in close proximity with helices α11 and α14 of NDC1-PD (Fig. 6*B*). In addition, the strands C and D of blade 3 (β3C & β3D) and the intervening 3AB loop from ALADIN may contact helix α16 of NDC1-PD.

The ALADIN-NDC1 hetero-dimer bridges the gap between Nup155-1 and Nup93-2 in the cytoplasmic half of the IR subunit (Fig. 6*C*). Specifically, the 1D2A loop and the 2CD loop of the ALADIN β-propeller are positioned close to the 1D2A loop and helix α3 of the N-terminal β-propeller of Nup155-1, respectively (Fig. 6*D*). Two surface loops from NDC1-PD, one between helices α12 and α13 and the other between α14 and α15, are located close to the surface loop between helices α10 and α11 of Nup93-2 (Fig. 6*E*).

The ALADIN-NDC1 hetero-dimer appears to play an important role in connecting Nup155-1 to the IR subunit. Nup155-1 and Nup155-1’, on the other hand, connect the IR to the CR and the NR, respectively (Fig. 5*A,B*). Any mutation in ALADIN that compromises its structural stability or its interaction with NDC1 is likely to have a negative impact on the connection of the IR with the CR/NR, which may consequently damage assembly and function of the NPC.

### Formation of the IR

We first aligned the 4.4-Å reconstruction of the IR subunit onto the EM map of the IR at 22-Å resolution. This practice was repeated eight times to generate a complete reconstruction of the IR. We then projected the atomic coordinates of the IR subunit into the reconstruction for each of the eight IR subunits, generating a composite model for the entire IR scaffold (Fig. 7*A*). This composite model contains 240 nucleoporins: 48 copies of Nup155, 32 copies each of Nup93 and CNT, and 16 copies each of Nup188, Nup205, ALADIN, and NDC1. The IR components Nup98 and Nup35 remain to be identified.

**Fig. 7.**
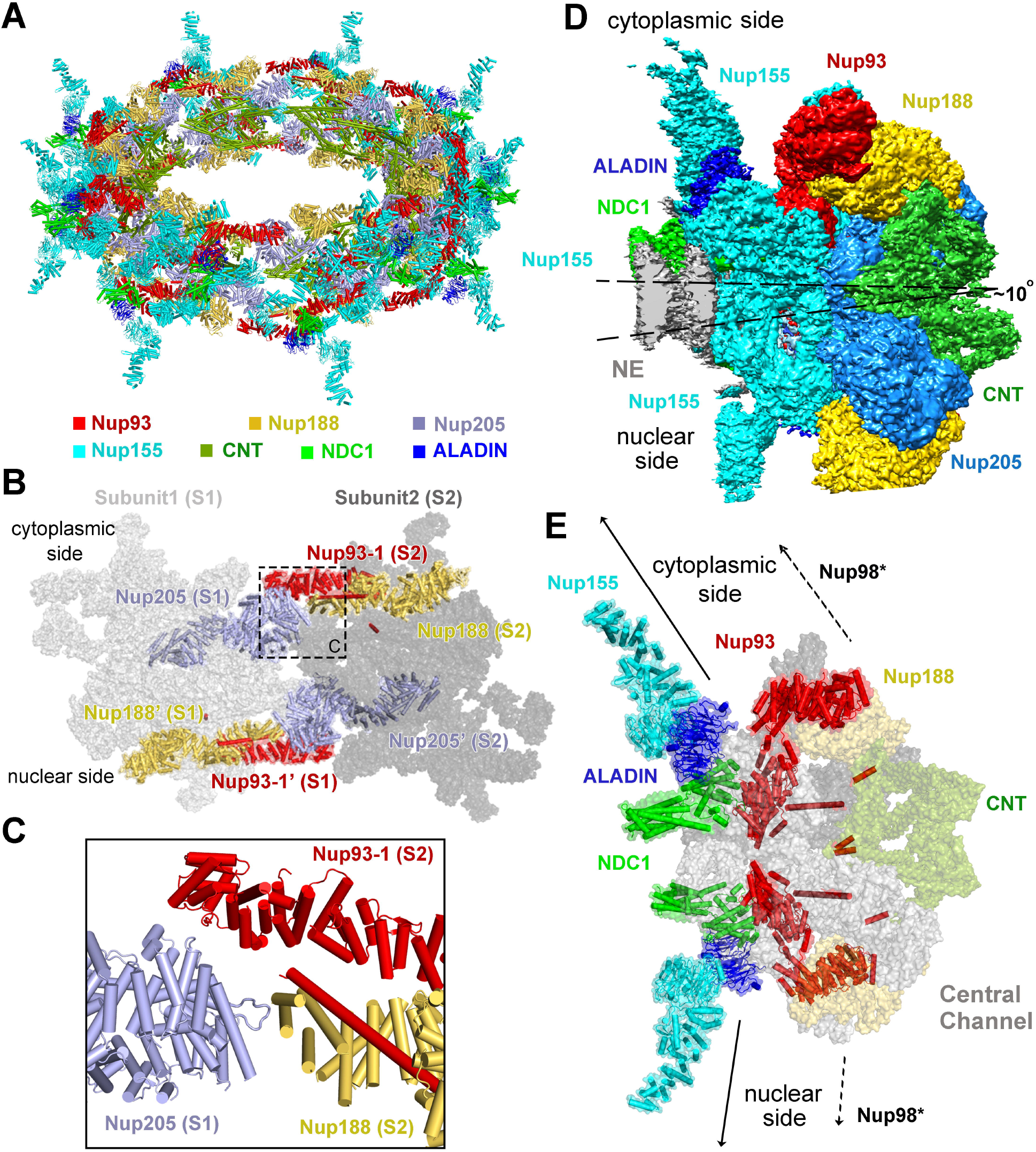
Structural features of the IR scaffold. (*A*) An overall view of the IR scaffold. (*B*) An overall view on inter-subunit interface between subunit 1 (S1) and subunit 2 (S2). (*C*) A close-up view on the cytoplasmic side of the interface between S1 and S2. Given the spatial gap in between, Nup205 of S1 may not directly interact with Nup93-1 or Nup188 from S2. (*D*) The IR subunit is slightly tilted towards the cytoplasmic side. (*E*) The IR subunit is linked to the outer rings through the linker nucleoporins Nup93, NDC1, ALADIN, and Nup155-1. The IR subunit is connected in opposite directions to the outer rings. Such connection depends on the ALADIN-NDC1 hetero-dimer to fill up the vacancy between Nup155-1 and Nup93-2.

The inter-subunit connection in the IR scaffold mainly involves Nup205 from subunit 1 (S1) and Nup93-1 and Nup188 from subunit 2 (S2) (Fig. 7*B*). The N-terminal helices of Nup205 from S1 are located close to the N-terminal helices of Nup93-1 and the C-terminal helices of Nup188 from S2 (Fig. 7*C*). Notably, however, the spatial gap between these potentially interacting helices from S1 and S2 is large enough to disallows direct interactions. This analysis suggests a weak interface between neighboring IR subunits, which is consistent with the observed contraction and dilation of the NPC pore (*20-22*). The small and large spatial gaps between two neighboring IR subunits may correspond to the contracted and dilated states of the NPC pore.

Unlike CR or NR, the IR has a nearly perfect internal symmetry (Fig. S15). However, the symmetric IR subunit is placed on the NE asymmetrically, tilting towards the CR side by approximately 10 degrees (Fig. 7*D*). The IR subunit is connected to the CR/NR through the linker nucleoporins Nup155, ALADIN, NDC1, and Nup93 (Fig. 7*E*). Such connection depends on the ALADIN-NDC1 hetero-dimer, which fills up the spatial gap between Nup155 and Nup93.

## Discussion

Advent of the atomic model of vertebrate IR subunit may serve as a framework for understanding the functional mechanism of NPC. A number of the nucleoporins are regulated by phosphorylation, which impacts on the assembly and function of NPC (*23, 24*). We mapped the phosphorylated sites onto the structure (*25, 26*) (Fig. S18). Ser307 and Ser675 of Nup155 face the nuclear membrane; phosphorylation of either residue is likely to alter their interaction with the membrane, thus potentially affecting membrane localization of Nup155 (Fig. S18*A)*. Phosphorylation of Ser117 in helix α5 of Nup93 may affect its interaction with Nup205 or Nup188; phosphorylation of Thr368 or Ser655 of Nup93 may alter the interface between Nup93-1 and Nup93-2 (Fig. S18*B)*. Analogously, phosphorylation of Nup188, Nup205 and CNT may change their local conformation or local environment for interaction (Fig. S18*C)*.

ALADIN is regulated by phosphorylation and is frequently targeted for mutation in the triple A syndrome (AAAS) (*18, 19*). All three potential phosphorylation sites (Ser186, Thr295, and Ser300 (*25, 26*)) in ALADIN are located on surface loops on the top side of the β-propeller (Fig. S19*A)*; these residues are likely involved in protein recruitment or interaction that might be sabotaged by mutations. Consistent with this analysis, this side of the ALADIN β-propeller is targeted by 15 AAAS-derived mutations (*27-40*) (Fig. S19*B)*. Presumably, these mutations either alter the structure or damage the interaction of ALADIN with other proteins. Other nucleoporins have been found to be targeted for mutations in other debilitating diseases, including Nup93 and Nup205 in steroid-resistant nephrotic syndrome (SRNS) (*41*) and Nup62 in infantile bilateral striatal necrosis (IBSN) (*42*) (Fig. S20*A,B)*. For example, the mutation Q416P, involving replacement of Gln416 by a helix-breaking residue Pro in the middle of an extended helix of Nup62, may destabilize its helical conformation (Fig. S20*B)*.

As a component of the central scaffold, Nup93 plays a key role in organizing the IR subunit through direct interactions with multiple nucleoporins (*16, 43*). Most notably, the Nup93 α-solenoid has an extended N-terminal extension that harbors two protein-interacting motifs (Fig. S5). The first motif is a short α-helix at the N-terminus of Nup93; it is involved in recruitment of the CNT (*12*). The second motif is an extended helix α5, which is known to interact with Nup205 or Nup188 (*12, 15*); it binds the axial groove of the C-terminal α-solenoid of Nup205 or Nup188. Remarkably, the interaction between Nup93-α5 and Nup205 axial groove in the IR subunit is exactly preserved in the CR subunit. A close examination of the Nup93-Nup205 interactions reveals an RMSD of no more than 4 Å over more than 1,200 aligned Cα atoms between the IR and CR subunits (Fig. S21).

The fact that the same helix α5 of Nup93 interacts similarly with Nup205 or Nup188 in both the CR and IR subunits suggests remarkable adaptability of Nup93. Indeed, the inherent conformational flexibility of Nup93 allows it to bridge interactions with a number of protein components in both CR and IR subunits. Such conformational flexibility is a hallmark of several other nucleoporins, as exemplified by Nup155 (Fig. S22). Although the two corresponding molecules of Nup155 in the cytoplasmic and nuclear halves are structurally identical (Fig. S22*A)*, different Nup155 molecules on the CR side exhibit very large RMSD values (Fig. S22*B)*.

Despite improvement in the quality of EM reconstruction for the vertebrate IR subunit, the average resolution of our EM reconstruction falls just short of the atomic range. The local resolution drops off quickly in the peripheral regions of the IR subunit. Consequently, we have been unable to identify the RRM-containing nucleoporin Nup35 or the linker nucleoporin Nup98. The current EM reconstruction suggests potential locations of Nup35 (Fig. S23). Additional improvement of the resolution should allow definitive assignment of Nup35 and Nup98.

In summary, we have determined the cryo-EM reconstruction of the *X. laevis* IR subunit at an average resolution of 4.4 Å, the highest achieved for the IR subunit of a vertebrate NPC. Structural analysis reveals a wealth of structural features that are likely to be functionally important.

## Acknowledgements

We thank the Cryo-EM Core and the Computing Core of Westlake University for technical support.

## Author Contributions

X.Zhu., P.W., and C. Z. prepared the sample. G.H., X.Zhan., C.Z., K.L. and J.L. collected the EM data. G.H., X.Zhan., C.Z., and K.L. processed the EM data. G.H. performed the cryo-EM SPA calculation. X.Zhan. built the atomic models. Q.Z., C.Y., Q.T. and M.L. provided critical support. All authors analyzed the structure. G.H., X.Zhan., C.Z., X.Zhu. and Y.S. wrote the manuscript. Y.S. conceived and supervised the project.

## Author Information

The authors declare no competing financial interests. Correspondence and requests for materials should be addressed to Y.S.

## Data and materials availability

The atomic coordinates of the IR subunit have been deposited in the Protein Data Bank with the accession code XXX. The EM map for the IR subunit has been deposited in the EMDB with the accession codes EMD-XXX.

## References and notes

1. C. Strambio-De-Castillia, M. Niepel, M. P. Rout, The nuclear pore complex: bridging nuclear transport and gene regulation. Nat Rev Mol Cell Bio 11, 490–501 (2010).

2. M. Beck, E. Hurt, The nuclear pore complex: understanding its function through structural insight. Nat Rev Mol Cell Bio 18, 73–89 (2017).

3. T. U. Schwartz, The Structure Inventory of the Nuclear Pore Complex. J Mol Biol 428, 1986–2000 (2016).

4. D. H. Lin, A. Hoelz, The Structure of the Nuclear Pore Complex (An Update). Annu Rev Biochem 88, 725–783 (2019).

5. Y. Zhang et al., Molecular architecture of the luminal ring of the Xenopus laevis nuclear pore complex. Cell Res 30, 532–540 (2020).

6. A. Hoelz, E. W. Debler, G. Blobel, The structure of the nuclear pore complex. Annu Rev Biochem 80, 613–643 (2011).

7. A. von Appen, M. Beck, Structure Determination of the Nuclear Pore Complex with Three-Dimensional Cryo electron Microscopy. J Mol Biol 428, 2001–2010 (2016).

8. J. Kosinski et al., Molecular architecture of the inner ring scaffold of the human nuclear pore complex. Science 352, 363–365 (2016).

9. G. Huang et al., Structure of the cytoplasmic ring of the Xenopus laevis nuclear pore complex by cryo-electron microscopy single particle analysis. Cell Res 30, 520–531 (2020).

10. X. Zhu et al., Near-atomic Structure of the Cytoplasmic Ring of the Xenopus laevis Nuclear Pore Complex. Submitted, (2021).

11. J. Jumper et al., Highly accurate protein structure prediction with AlphaFold. Nature 596, 583–589 (2021).

12. T. Stuwe et al., Architecture of the fungal nuclear pore inner ring complex. Science 350, 56–64 (2015).

13. B. Kind, K. Koehler, M. Lorenz, A. Huebner, The nuclear pore complex protein ALADIN is anchored via NDC1 but not via POM121 and GP210 in the nuclear envelope. Biochem Biophys Res Commun 390, 205–210 (2009).

14. Y. Yamazumi et al., The transmembrane nucleoporin NDC1 is required for targeting of ALADIN to nuclear pore complexes. Biochem Biophys Res Commun 389, 100–104 (2009).

15. D. H. Lin et al., Architecture of the symmetric core of the nuclear pore. Science 352, aaf1015 (2016).

16. R. Sachdev, C. Sieverding, M. Flotenmeyer, W. Antonin, The C-terminal domain of Nup93 is essential for assembly of the structural backbone of nuclear pore complexes. Mol Biol Cell 23, 740–749 (2012).

17. C. Bizzarri, D. Benevento, C. Terzi, A. Huebner, M. Cappa, Triple A (Allgrove) syndrome: an unusual association with syringomyelia. Ital J Pediatr 39, 39 (2013).

18. A. Huebner et al., The triple A syndrome is due to mutations in ALADIN, a novel member of the nuclear pore complex. Endocr Res 30, 891–899 (2004).

19. K. Handschug et al., Triple A syndrome is caused by mutations in AAAS, a new WD-repeat protein gene. Hum Mol Genet 10, 283–290 (2001).

20. C. E. Zimmerli et al., Nuclear pores constrict upon energy depletion. bioRxiv, 2020.2007.2030.228585 (2020).

21. T. R. Lezon, A. Sali, I. Bahar, Global motions of the nuclear pore complex: insights from elastic network models. PLoS Comput Biol 5, e1000496 (2009).

22. A. P. Schuller et al., The cellular environment shapes the nuclear pore complex architecture. Nature 598, 667–671 (2021).

23. J. S. Glavy et al., Cell-cycle-dependent phosphorylation of the nuclear pore Nup107-160 subcomplex. P Natl Acad Sci USA 104, 3811–3816 (2007).

24. F. Huguet, S. Flynn, P. Vagnarelli, The Role of Phosphatases in Nuclear Envelope Disassembly and Reassembly and Their Relevance to Pathologies. Cells-Basel 8, (2019).

25. N. Dephoure et al., A quantitative atlas of mitotic phosphorylation. P Natl Acad Sci USA 105, 10762–10767 (2008).

26. J. V. Olsen et al., Quantitative phosphoproteomics reveals widespread full phosphorylation site occupancy during mitosis. Sci Signal 3, ra3 (2010).

27. M. Hirano, Y. Furiya, H. Asai, A. Yasui, S. Ueno, ALADINI482S causes selective failure of nuclear protein import and hypersensitivity to oxidative stress in triple A syndrome. P Natl Acad Sci USA 103, 2298–2303 (2006).

28. M. Krumbholz, K. Koehler, A. Huebner, Cellular localization of 17 natural mutant variants of ALADIN protein in triple A syndrome - shedding light on an unexpected splice mutation. Biochem Cell Biol 84, 243–249 (2006).

29. K. Koehler et al., Axonal neuropathy with unusual pattern of amyotrophy and alacrima associated with a novel AAAS mutation p.Leu430Phe. Eur J Hum Genet 16, 1499–1506 (2008).

30. A. Salmaggi et al., Late-onset triple A syndrome: a risk of overlooked or delayed diagnosis and management. Horm Res 70, 364–372 (2008).

31. C. Villanueva-Mendoza, O. artinez-Guzman, D. Rivera-Parra, J. C. Zenteno, Triple A or Allgrove syndrome. A case report with ophthalmic abnormalities and a novel mutation in the AAAS gene. Ophthalmic Genet 30, 45–49 (2009).

32. M. Luigetti et al., Triple A syndrome: a novel compound heterozygous mutation in the AAAS gene in an Italian patient without adrenal insufficiency. J Neurol Sci 290, 150–152 (2010).

33. T. Milenkovic et al., Triple A syndrome: 32 years experience of a single centre (1977-2008). Eur J Pediatr 169, 1323–1328 (2010).

34. K. Nakamura et al., Adult or late-onset triple A syndrome: case report and literature review. J Neurol Sci 297, 85–88 (2010).

35. A. Dixit, G. Chow, A. Sarkar, Neurologic presentation of triple A syndrome. Pediatr Neurol 45, 347–349 (2011).

36. M. Dumic et al., Two siblings with triple A syndrome and novel mutation presenting as hereditary polyneuropathy. Eur J Pediatr 170, 393–396 (2011).

37. A. E. Vallet et al., Neurological features in adult Triple-A (Allgrove) syndrome. J Neurol 259, 39–46 (2012).

38. M. Capataz Ledesma, P. Mendez Perez, R. Rodriguez Lopez, E. Galan Gomez, [Allgrove syndrome (triple A). Finding of a mutation not described in the AAAS gene]. An Pediatr (Barc) 78, 109–112 (2013).

39. L. Papageorgiou, K. Mimidis, K. R. Katsani, G. Fakis, The genetic basis of triple A (Allgrove) syndrome in a Greek family. Gene 512, 505–509 (2013).

40. F. Roucher-Boulez et al., Triple-A syndrome: a wide spectrum of adrenal dysfunction. Eur J Endocrinol 178, 199–207 (2018).

41. D. A. Braun et al., Mutations in nuclear pore genes NUP93, NUP205 and XPO5 cause steroid-resistant nephrotic syndrome. Nat Genet 48, 457–465 (2016).

42. L. Basel-Vanagaite et al., Mutated nup62 causes autosomal recessive infantile bilateral striatal necrosis. Ann Neurol 60, 214–222 (2006).

43. B. Vollmer, W. Antonin, The diverse roles of the Nup93/Nic96 complex proteins - structural scaffolds of the nuclear pore complex with additional cellular functions. Biol Chem 395, 515–528 (2014).

