## Supplemental Information for "Cryo-EM structure of the Inner Ring from *Xenopus laevis* Nuclear Pore Complex"

### METHODS

#### Cryo-EM sample preparation and EM data acquisition

The same cryo-EM data sets used for reconstruction of the cytoplasmic ring (CR) subunit from *Xenopus laevis* (*X. laevis*) oocytes (*1*) were used for reconstruction of the inner ring (IR) subunit. In short, the nuclear envelope (NE) from *X. laevis* oocytes was prepared as described (*2, 3*). The cryo-EM sample was prepared as described (*1*). The gold EM grids (R1.2/1.3&R2/1&R2/2; Quantifoil, Jena, Germany) were blotted for 8 seconds with a blot force of 15 and vitrified by plunge-freezing into liquid ethane using a Vitrobot Mark IV (Thermo Fisher Scientific) at 8 °C under 100% humidity.

Details of data acquisition are as described (*1*). With the grids tilting at angles of 0, 30, 45, and 55 degrees, 46,143 micrographs were recorded on a Titan Krios electron microscope (FEI) operating at 300 kV with a nominal magnification of 64,000x and equipped with a Gatan GIF Quantum energy filter (slit width 20 eV) (Table S1). A K3 detector (Gatan Company) was used in the super-resolution mode, with a calibrated pixel size of 0.6935 Å for the movie files (Table S1). The movie images were binned twice during motion correction, arriving at a pixel size of 1.387 Å for the final motion corrected images. All frames in each stack were first aligned and summed using MotionCor2 (*4*). Dose weighting was performed using MotionCor2 (*4*). The average defocus values were set between -1.5 and -3.0 μm and were estimated using Gctf (*5*).

**An initial model of the IR from *X. laevis* oocytes**

33,747 micrographs were manually selected from the original dataset of 46,143 micrographs. A total of 800,825 particles were manually selected from these micrographs (Fig. S1*A*). Initial defocus estimation was carried out as previously described (*3*) prior to all other data processing procedures.

The central portion of an NPC comprises four ring scaffolds: CR, IR, nuclear ring (NR), and luminal ring (LR). Due to the inherent flexibility among the four rings, it is practically impossible to refine the entire NPC as a single particle to high resolution. We therefore carried out initial pose estimation of the NPC particles on one of the relatively stable ring scaffolds: the CR. The NPC particles were first aligned to the CR side as described (*1*). Following refinement of the CR structure, we continued the 3D refinement procedure from the last iteration with a layered mask focusing on the IR and LR layer. Only pixels within a specific layer, with z_start < z < z_end, have pixel values of 1; all other pixels that have z coordinates below or above this layer have zero values. Transition between the two regions follows a raised cosine scheme. The continue refinement yielded a reconstruction at 22 Å resolution based on 660,302 NPC particles (Fig. S1*A*).

The resulting data star files from the final round of auto-refinement were imported into cryoSparc (*6*) for a round of Local-Refinement to arrive at a reconstruction at 22 Å resolution (Fig. S1*A*). The nominal resolution is the same as that reported in RELION (*7*) but the EM map has significantly less noise in the CR region (Fig. S1*A*). The C8 symmetry was applied throughout this stage of data processing. The IR from *X. laevis* exhibits an inner diameter of 45 nm, close to that reported in human NPC from purified NE (*8*) (Fig. S1*B*). But this inner diameter is significantly constricted when compared to that reported in the *in situ* structures of human NPC (*9, 10*).

**Data processing and reconstruction of the IR subunit**

We extracted the IR subunit particles based on the alignment parameters of the 22-Å IR reconstruction. We updated the orientation, shift and defocus parameters for each subunit according to a published protocol (*3*). 5,223,773 particles of the IR subunit were extracted using a box size of 128 and a binned pixel size of 5.548 Å (Fig. S2*A*). We performed one round of 3D classification (K=1) with 10 iterations. The data star file from iteration 10 was then used for re-extraction of bin2 particles with a box size of 256 and a binned pixel size of 2.774 Å (Fig. S2*A*). The entire dataset of bin2 particle of the IR subunit were then subjected to three rounds of parameter refinement (*1*). This practice allowed selection of 2,139,754 particles, which yielded a reconstruction of the IR subunit at 5.6 Å resolution.

To fully utilize the dataset, these IR subunits were projected back to the original IR particles. The defocus values of all subunit particles within the same IR were pooled together to calculate a corrected average defocus value for the center of mass of each IR particle. All subunits of this IR were then re-extracted using the updated defocus value deduced from the corrected average defocus of this IR particle.

Through this procedure, 3,013,260 subunit particles were extracted, resulting in a reconstruction at 5.6 Å resolution after auto-refinement (Fig. S2*A*).

This data set was then used for re-extraction with a box size of 400 and pixel size of 1.387 Å (Fig. S2*B*). Data processing beyond this point was carried out as described (*3*). In short, the extracted particles were directly subjected to one round of CTF-refinement followed by one round of auto-refinement, resulting in a reconstruction of the IR subunit at 5.2 Å resolution (Fig. S2*B*). Three additional rounds of parameter refinement cycles were repeated to arrive at a final average resolution of 4.4 Å from 2,093,631 particles (Fig. S2*B*). The angular distribution of the particles is reasonable and the overall reconstruction has isotropic resolution (Fig. S3). The final EM map displays clear features for secondary structural elements (Figs. S4-S7). These features facilitated sequence assignment of the nucleoporins (Figs. S8-S13).

**Atomic modeling of the IR subunit**

The previous EM map of the IR subunit from human NPC, achieved through cryo-ET analysis, displays an average resolution of about 21 Å (*11*). The atomic coordinates of human IR (PDB: 5IJO) (*11*) were manually fitted into our 4.4-Å reconstruction of the *X. laevis* IR subunit using Chimera (*12*). The much improved EM density maps allowed unambiguous assignment of most IR components and accurate placement of secondary structural elements. This practice allows identification of 30 molecules of nucleoporins in each IR subunit, including four copies of Nup93, six copies of

Nup155, four channel nucleoporin heterotrimers (CNT, composed of Nup62, Nup58 and Nup54) (*13*), two copies of NDC1, two copies of ALADIN, two copies of Nup205 and two copies of Nup188.

The atomic coordinates of Nup93 and Nup205 from the CR subunit (*1*) were docked into the EM maps with little adjustment. Using the recently released structure prediction tool AlphaFold (*14*), we generated the atomic coordinates for each of the other *X. laevis* proteins. These predicted structures for most *X. laevis* nucleoporins are remarkably similar to the homology-modeled structures of the corresponding human nucleoporins. The predicted structure of the CNT (Nup62, Nup58 and Nup54) is nearly identical to that of the reported structure for *X. laevis* CNT (PDB: 5C3L) (*15*). We individually docked each predicted structure of the *X. laevis* nucleoporin into the 4.4-Å EM map and made manual adjustment using Coot (*16*). The final atomic model of the *X. laevis* IR subunit contains 19,325 amino acids, with 788 α-helices and 312 β-strands.

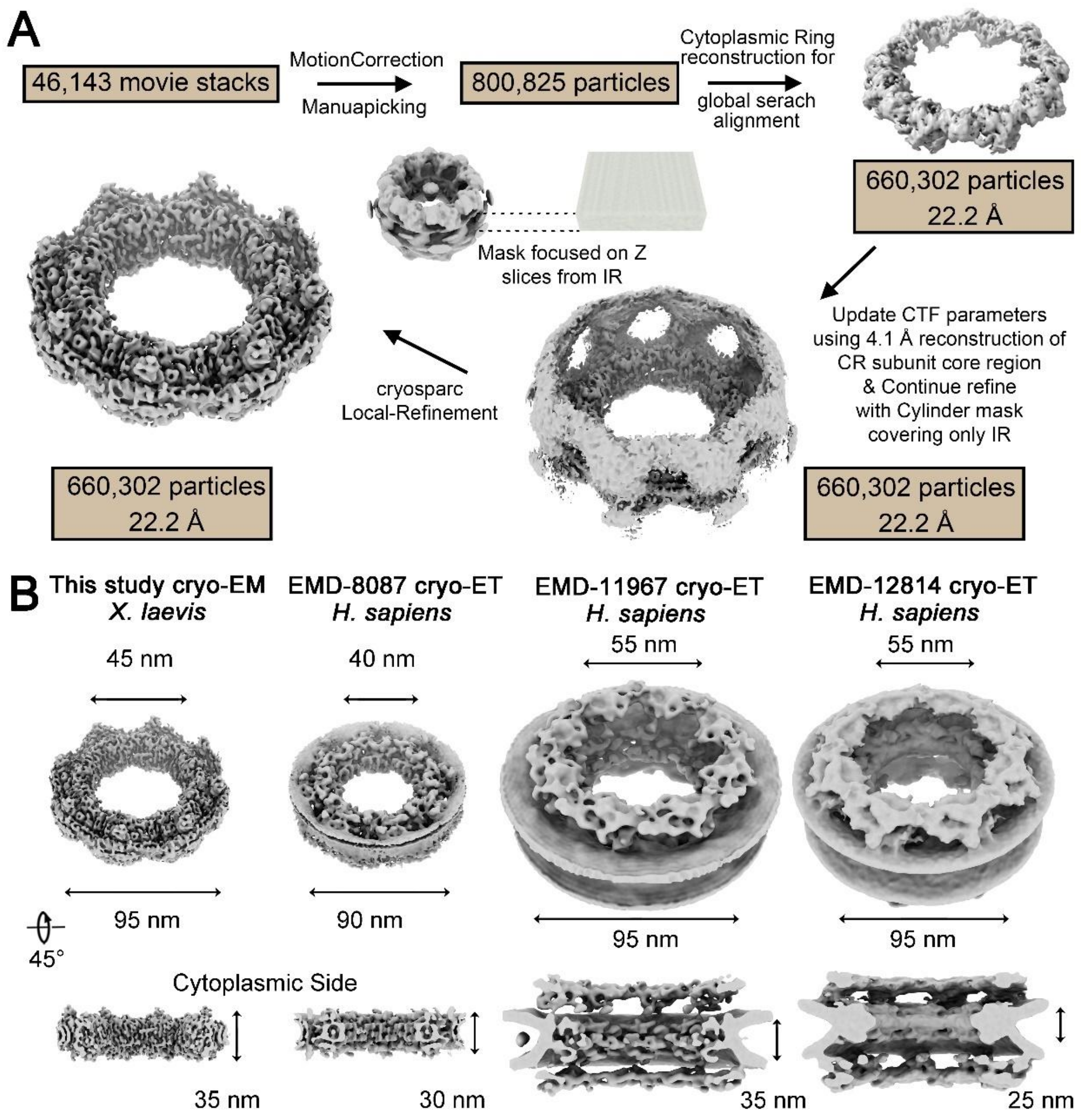


**Fig. S1 Cryo-EM analysis for the IR of NPC from *X. laevis* oocytes.** (*A*) A flowchart of data analysis for reconstruction of the IR from *X. laevis* NPC at 22 Å resolution. (*B*) Comparison of overall structural features between the IR of *X. laevis* NPC and IR of human NPC. Notably, the experimental methods for structure determination are also different.

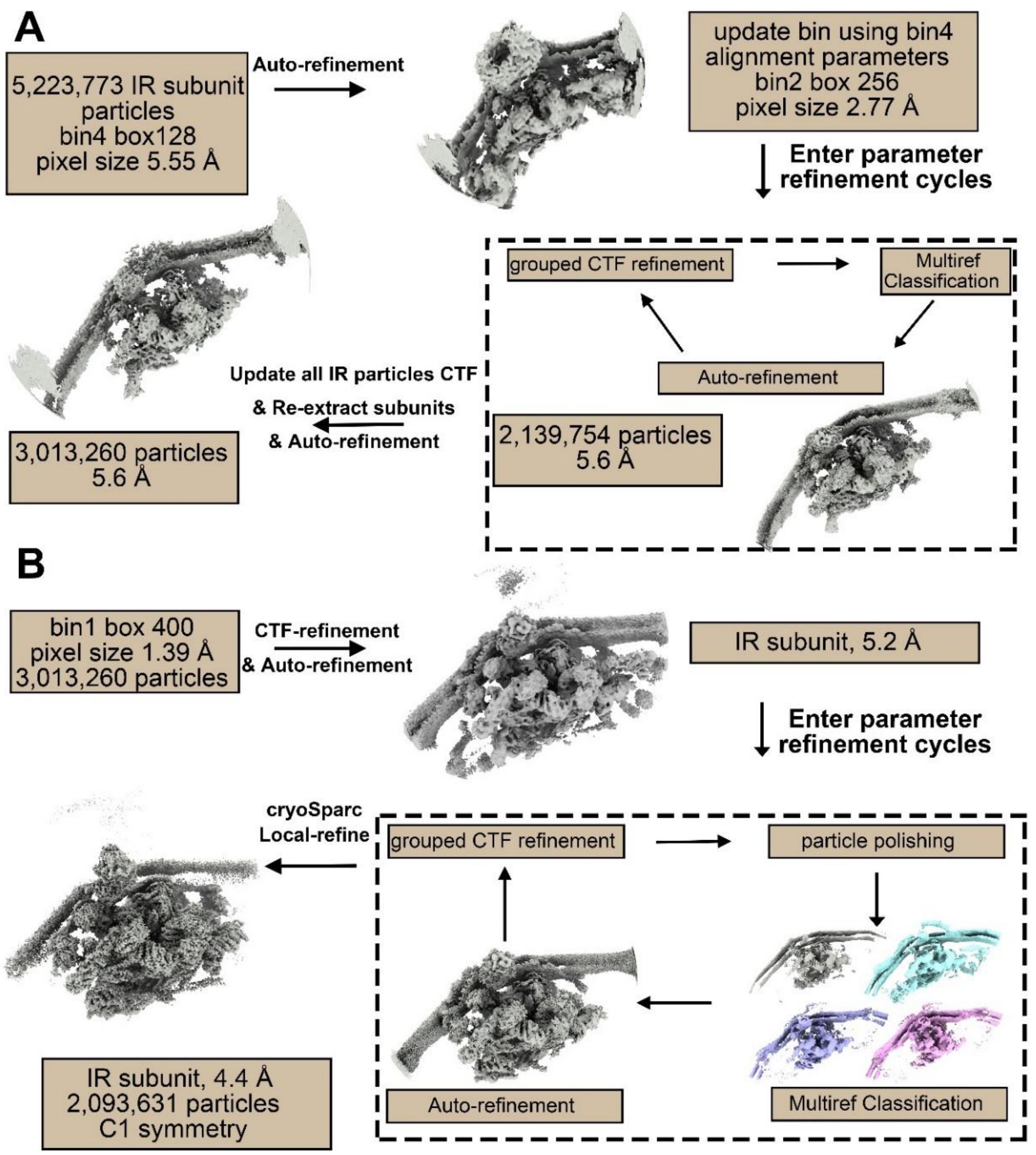


**Fig. S2 Processing of the cryo-EM data for reconstruction of the IR subunit from *X. laevis* NPC.** (*A*) A flowchart of data processing for reconstruction of the IR subunit to an average resolution of 5.6 Å. This part of the data analysis only involves the bin-4 and bin-2 levels. (*B*) A flowchart of data processing for reconstruction of the IR subunit to an average resolution of 4.4 Å. This part of the data analysis involves the bin-1 level. For details of either panel *A* or *B*, please refer to the section "Data processing and reconstruction of the IR subunit" in the Methods.

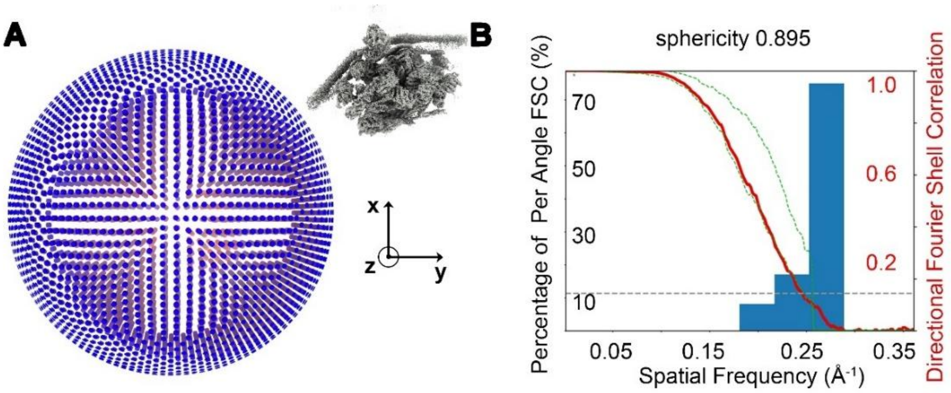


**Fig. S3 Cryo-EM analysis of the IR subunit.** (*A*) Angular distribution of the single-particle cryo-EM reconstruction for the IR subunit. Each cylinder represents one view and the height of the cylinder is proportional to the number of particles for that view. (*B*) Directional Fourier Shell correlation (FSC) curves and directional FSC histograms for cryo-EM reconstruction of the IR subunit. All directional FSC curves were calculated using the following website: https://3dfsc.salk.edu (*17*).

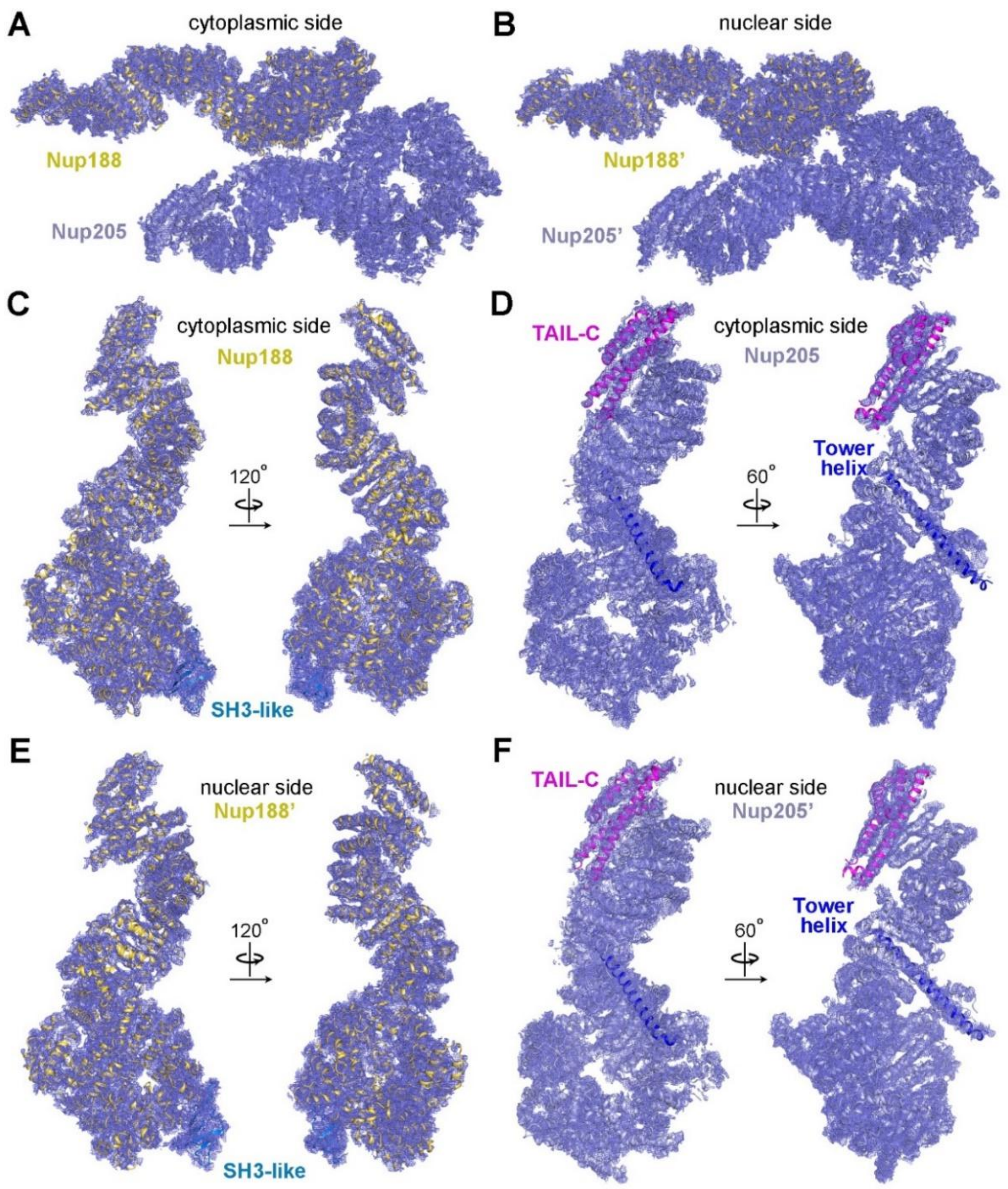


**Fig. S4 The EM density maps for Nup188 and Nup205.** (*A*) The overall EM density map of Nup188 and Nup205 from the cytoplasmic side. (*B*) The overall EM density map of Nup188 and Nup205 from the nuclear side (Nup188' and Nup205'). (*C*) The EM density map of Nup188 is displayed in two related views. The position of the SH3-like domain is indicated. (*D*) The EM density map of Nup205 is displayed in two related views. The positions of TAIL-C and Tower helix are indicated. (*E*) The EM density map of Nup188' is displayed in two related views. The position of the SH3-like domain is indicated. (*F*) The EM density map of Nup205' is displayed in two related views. The positions of TAIL-C and Tower helix are indicated. All EM density maps in this and following figures were prepared using the reconstruction of the IR subunit with a contour level between 5σ and 10σ.

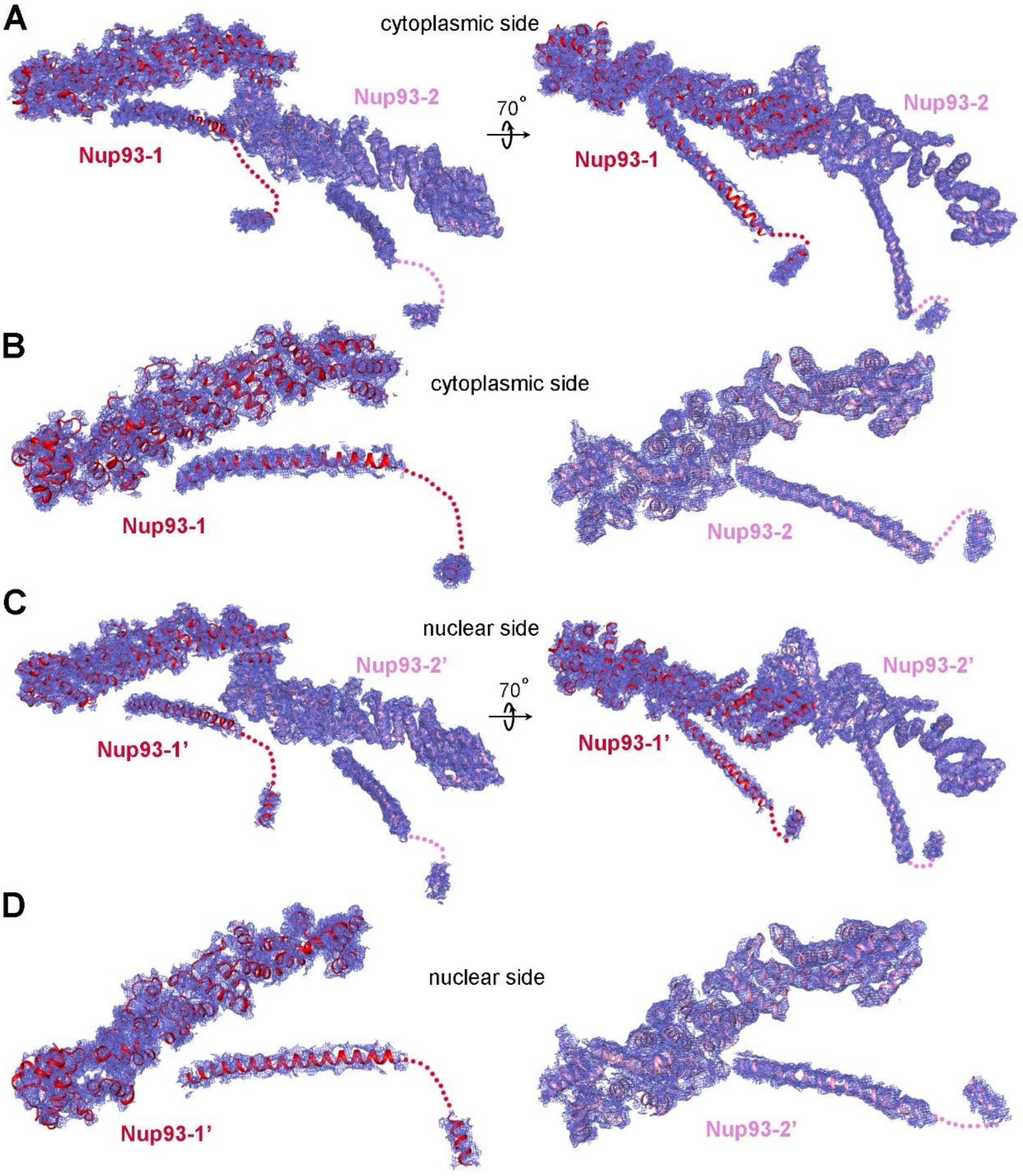


**Fig. S5 The EM density maps for Nup93.** (*A*) The overall EM density map of Nup93 on the cytoplasmic side. Two related views are shown. The two Nup93 molecules are designated Nup93-1 and Nup93-2, the latter of which is located closer to the center of the IR subunit. (*B*) The EM density maps of Nup93-1 (left panel) and Nup93-2 (right panel). (*C*) The overall EM density map of Nup93 on the nuclear side is shown in two related views. The two Nup93 molecules are designated Nup93-1' and Nup93-2', which correspond to Nup93-1 and Nup93-2 by C2 symmetry, respectively. (*D*) The EM density maps of Nup93-1' (left panel) and Nup93-2' (right panel).

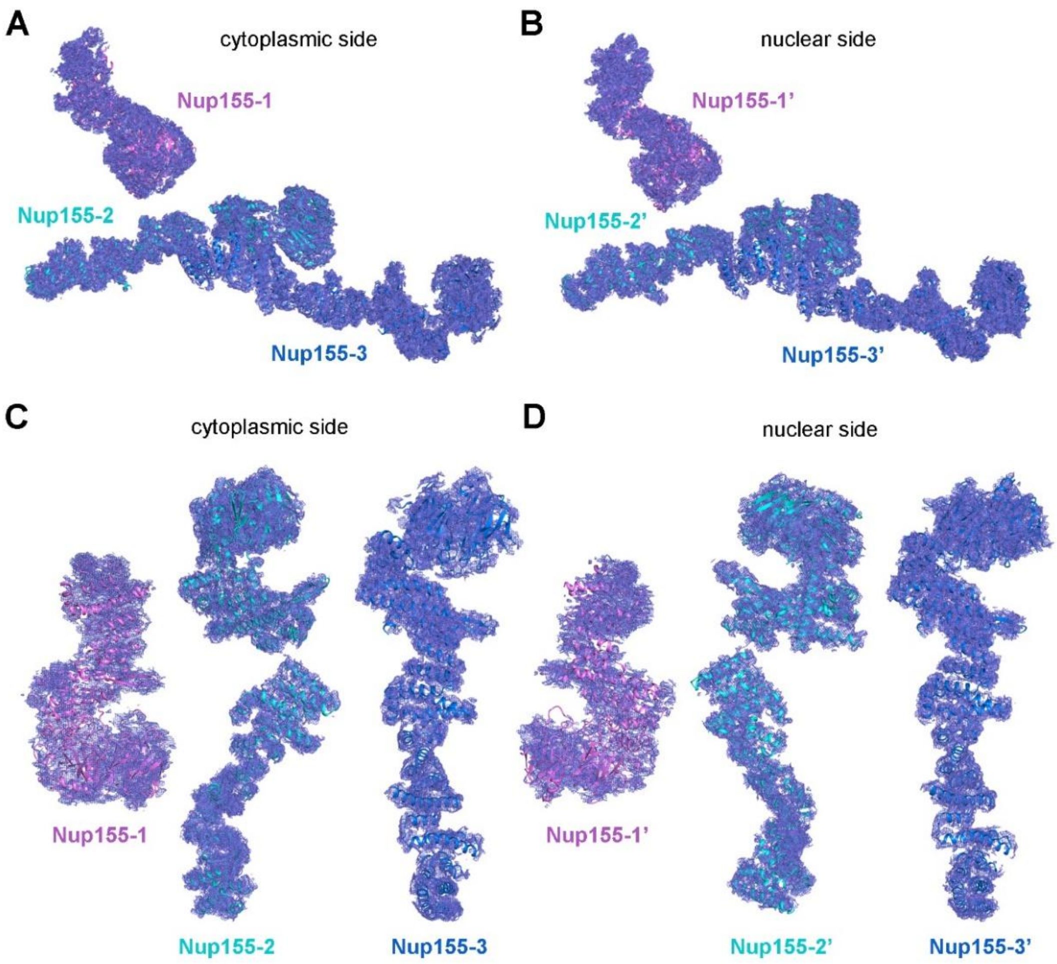


**Fig. S6 The EM density maps for Nup155.** (*A*) The overall EM density map of Nup155 on the cytoplasmic side. The three Nup155 molecules are designated Nup155-1, Nup155-2 and Nup155-3. (*B*) The overall EM density map of Nup155 on the nuclear side. The three Nup155 molecules are designated Nup155-1', Nup155-2' and Nup155-3', which correspond to Nup155-1, Nup155-2 and Nup155-3 by C2 symmetry, respectively. (*C*) The EM density maps of Nup155-1 (left panel), Nup155-2 (middle panel), and Nup155-3 (right panel). (*D*) The EM density maps of Nup155-1' (left panel), Nup155-2' (middle panel), and Nup155-3' (right panel).

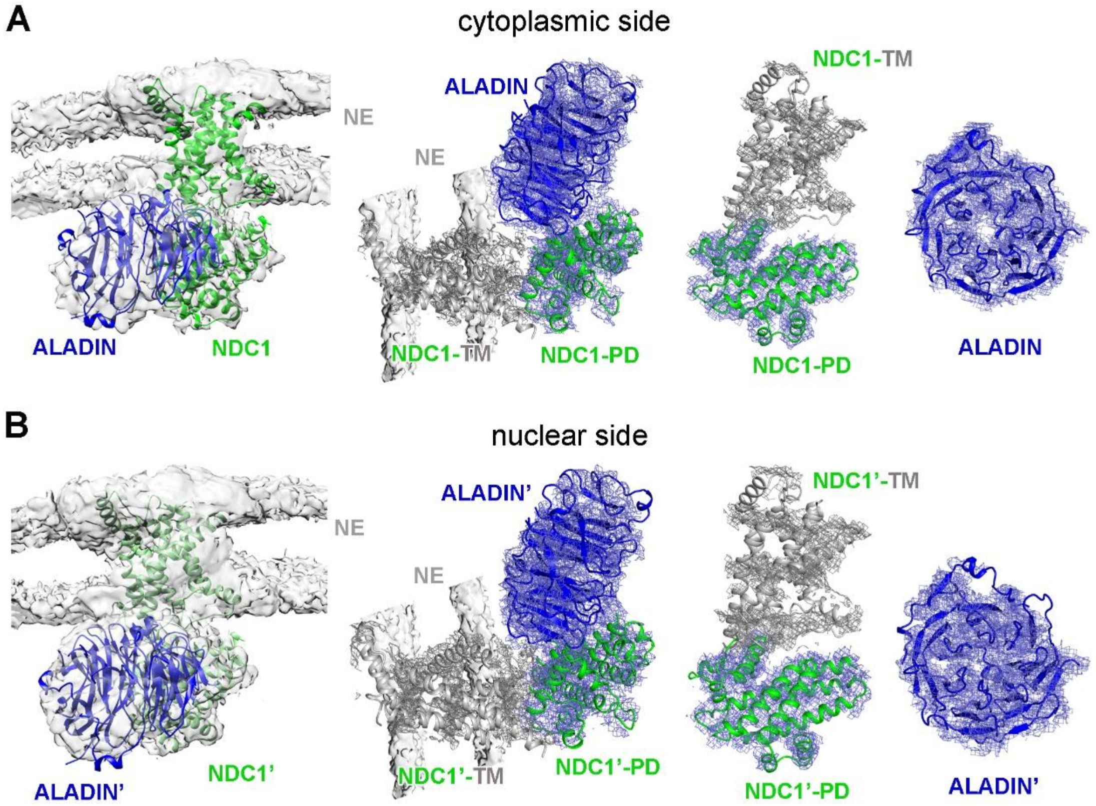


**Fig. S7 The EM density maps for ALADIN and NDC1.** (*A*) The EM density maps of ALADIN and NDC1 on the cytoplasmic side. Placement of ALADIN and NDC1 into the EM density maps is shown in two related views (two left panels). The individual EM maps for ALADIN and NDC1 are shown (two right panels). (*B*) The EM density maps of ALADIN and NDC1 on the nuclear side. ALADIN and NDC1 on the nuclear side are designated ALADIN' and NDC1', respectively. Placement of ALADIN' and NDC1' into the EM density maps is shown in two related views (two left panels). The individual EM maps for ALADIN' and NDC1' are shown (two right panels). PD, pore domain; TM, transmembrane domain.

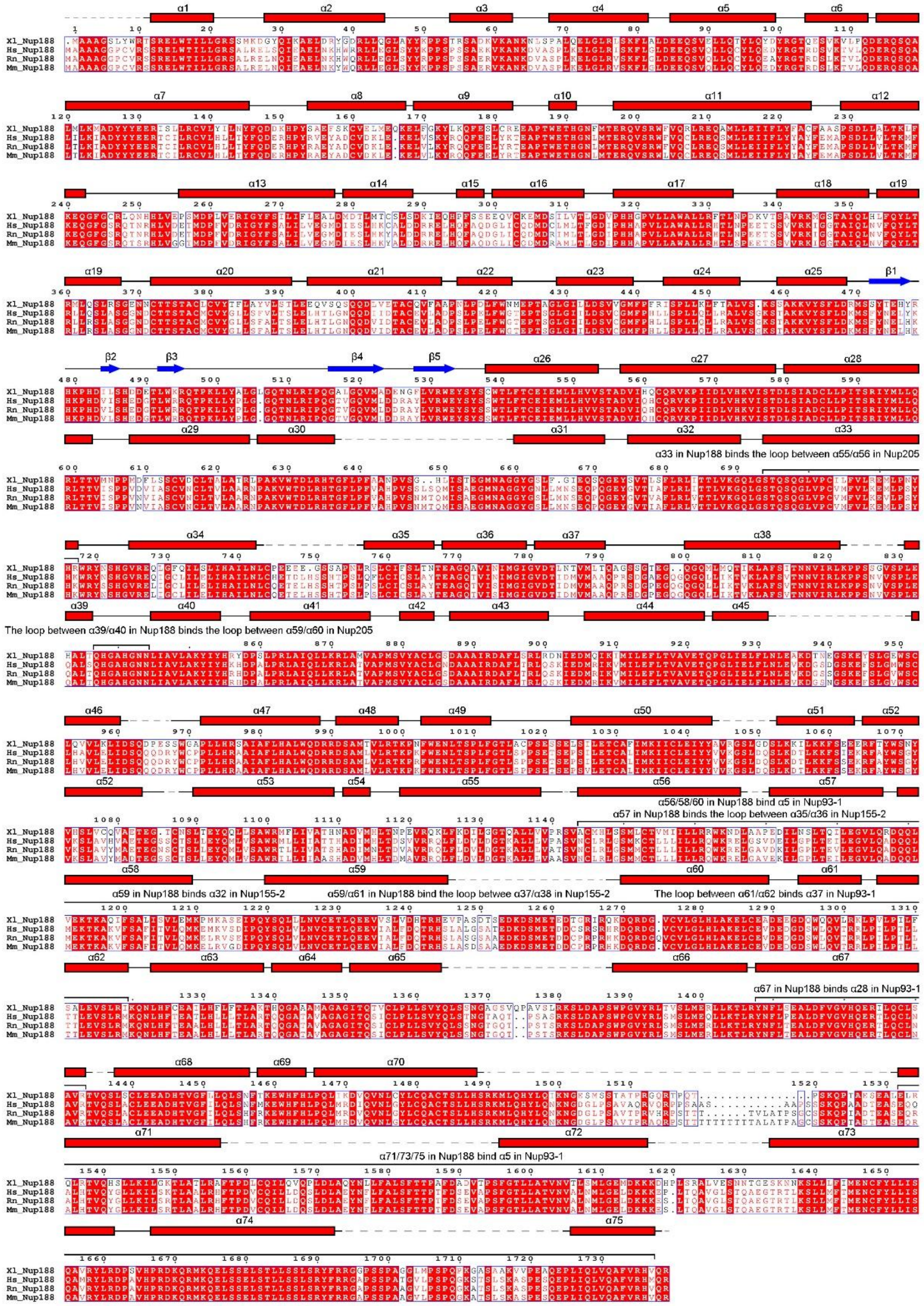


**Fig. S8 Sequence alignment among Nup188 orthologues from *X. laevis* (Xl), *Homo sapiens* (Hs), *Rattus norvegicus* (Rn), and *Mus musculus* (Mm).** The sequence alignment of the full-length Nup188 is shown. Conserved amino acids are boxed, with invariant residues highlighted in red background. The observed secondary structural elements in both Nup188 molecules are indicated above the sequences. Structural elements interacting with other nucleoporins in the IR subunit are indicated.

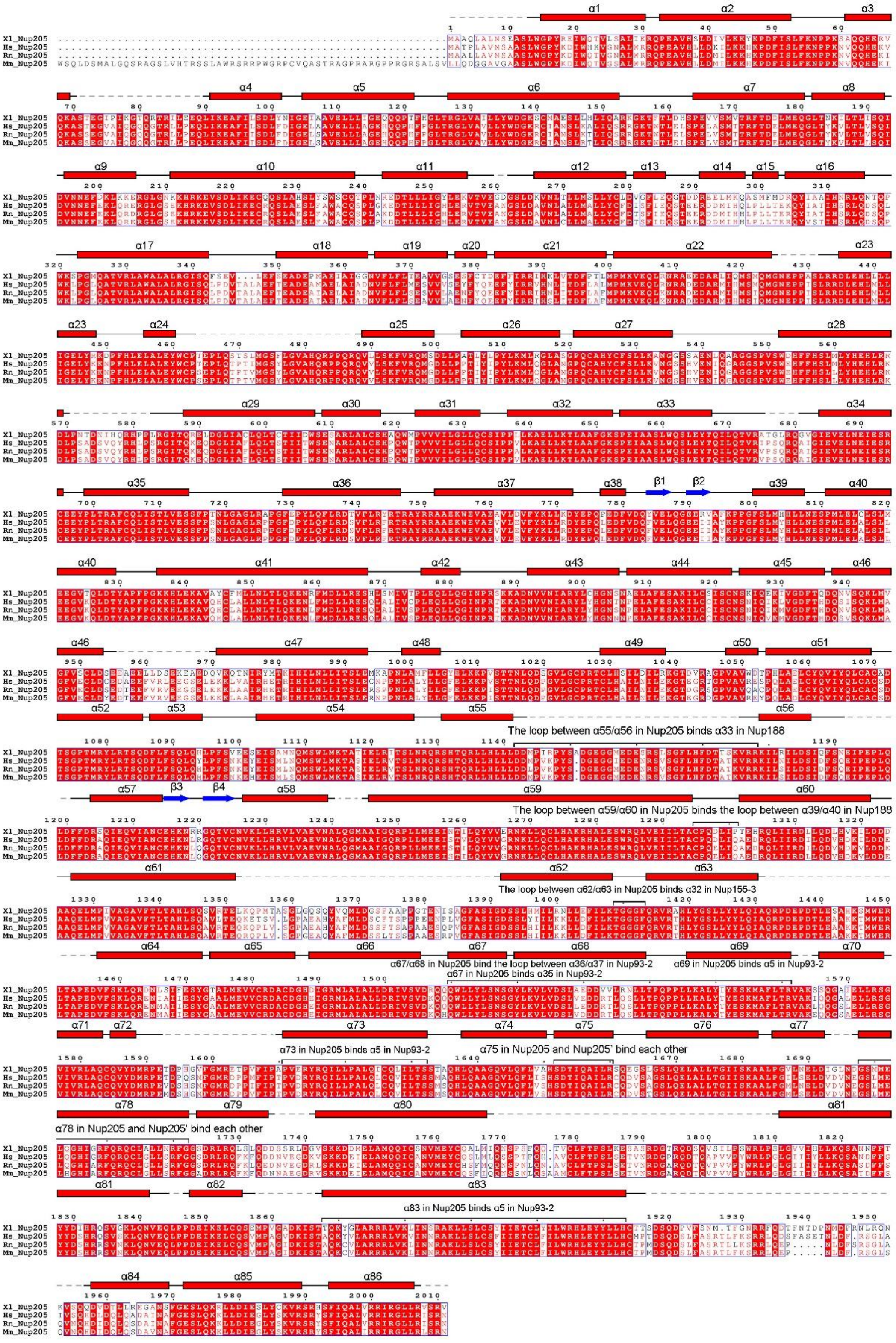


**Fig. S9 Sequence alignment among Nup205 orthologues from multiple vertebrates.** Conserved amino acids are boxed. Invariant residues are highlighted in red background. The observed secondary structural elements in both Nup205 molecules from IR subunit are indicated above the sequences. The structural elements interacting with other nucleoporins in the IR subunit are also indicated.

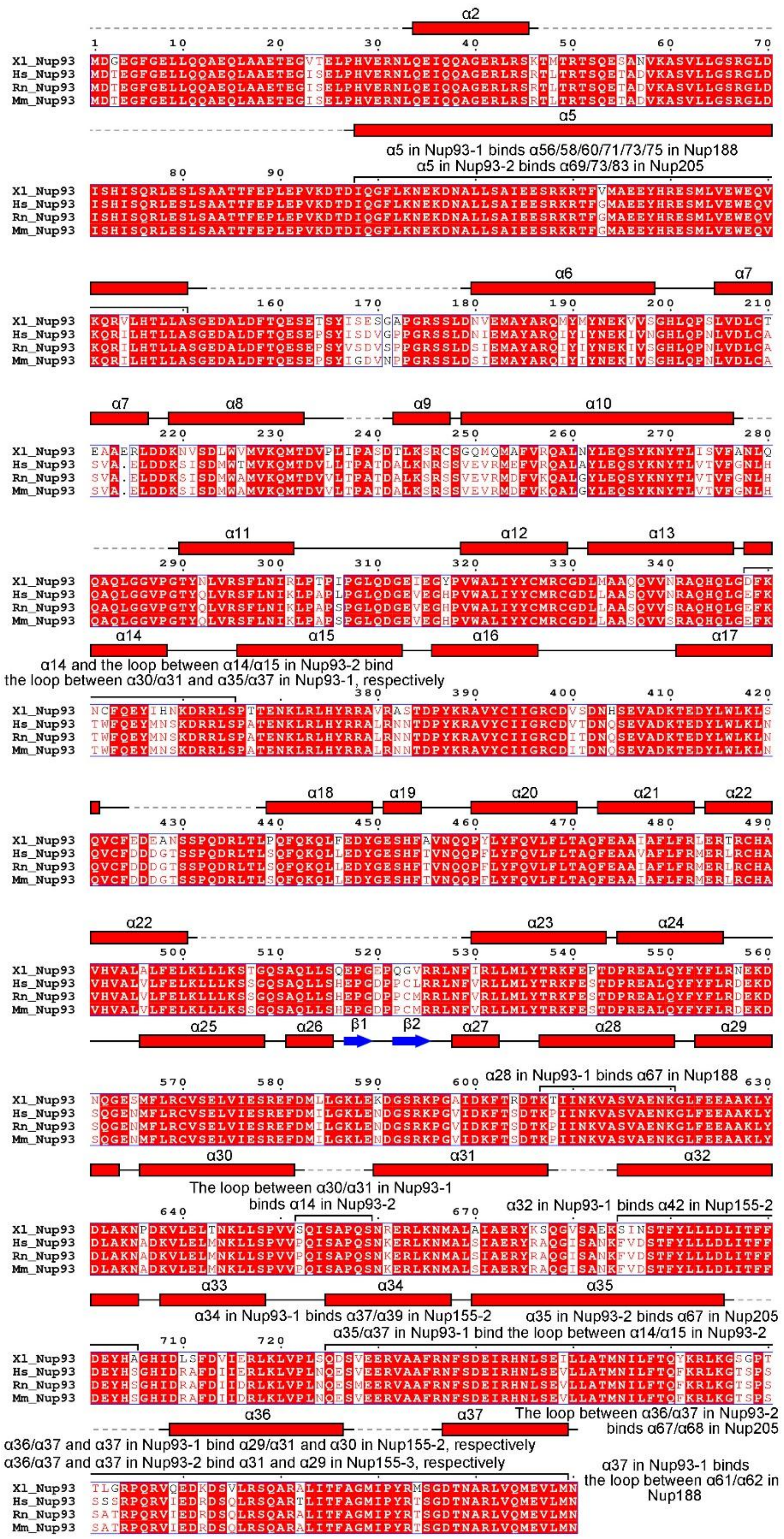


**Fig. S10 Sequence alignment among Nup93 orthologues from multiple vertebrates.** Conserved amino acids are boxed. Invariant residues are highlighted in red background. The observed secondary structural elements in all four Nup93 molecules are indicated above the sequences. The structural elements interacting with other nucleoporins in the IR subunit are also indicated.

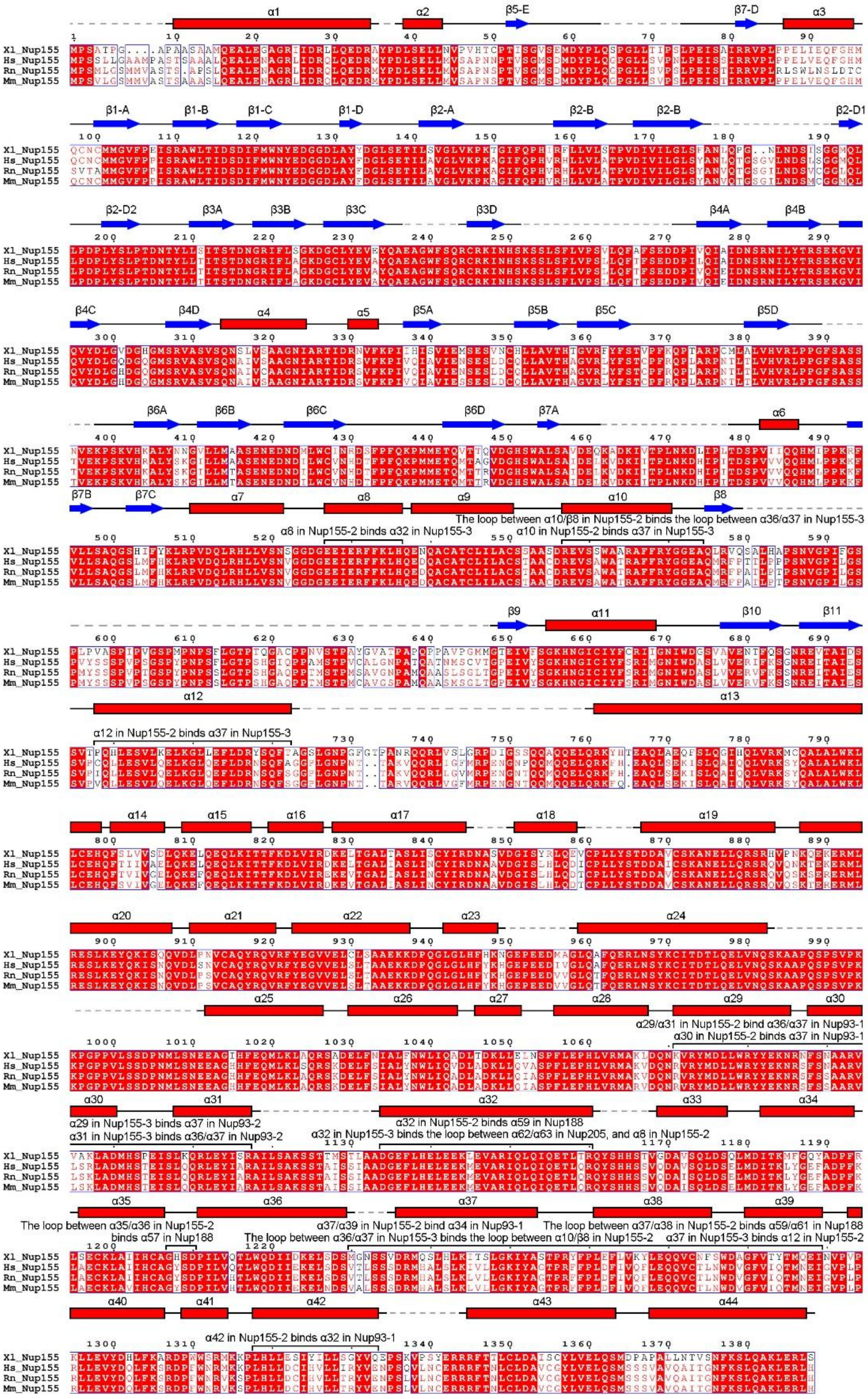

**Fig. S11 Sequence alignment among Nup155 orthologues from multiple vertebrates.** The conserved amino acids are boxed, with invariant residues highlighted in red background. The observed secondary structural elements in Nup155-2 and Nup155-3 are indicated above the sequences. The structural elements interacting with other nucleoporins in the IR subunit are also indicated.

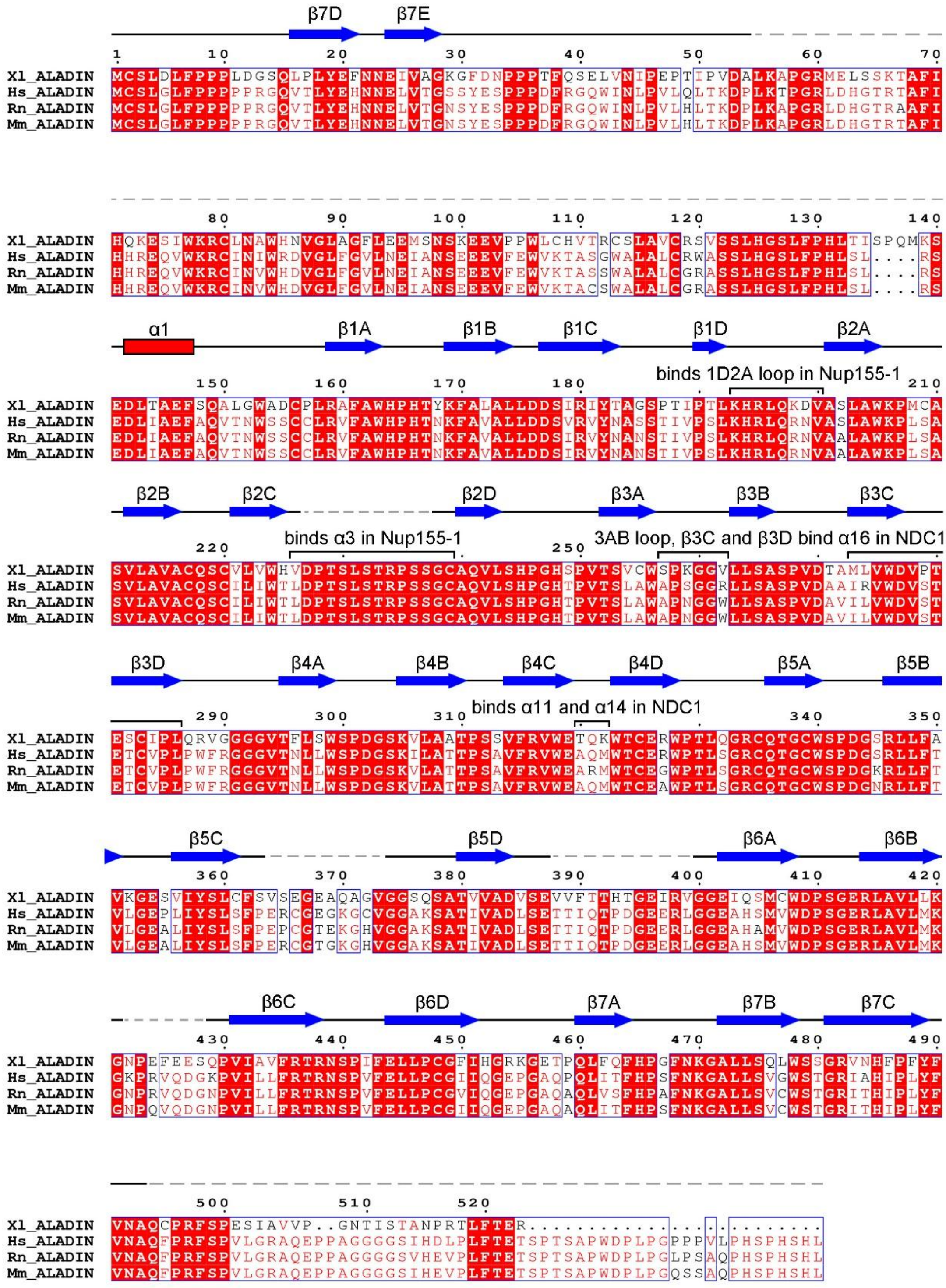


**Fig. S12 Sequence alignment among ALADIN orthologues from multiple vertebrates.** The conserved amino acids are boxed, with invariant residues highlighted in red background. The observed secondary structural elements in both ALADIN molecules are indicated above the sequences. The structural elements interacting with other nucleoporins in the IR subunit are also indicated.

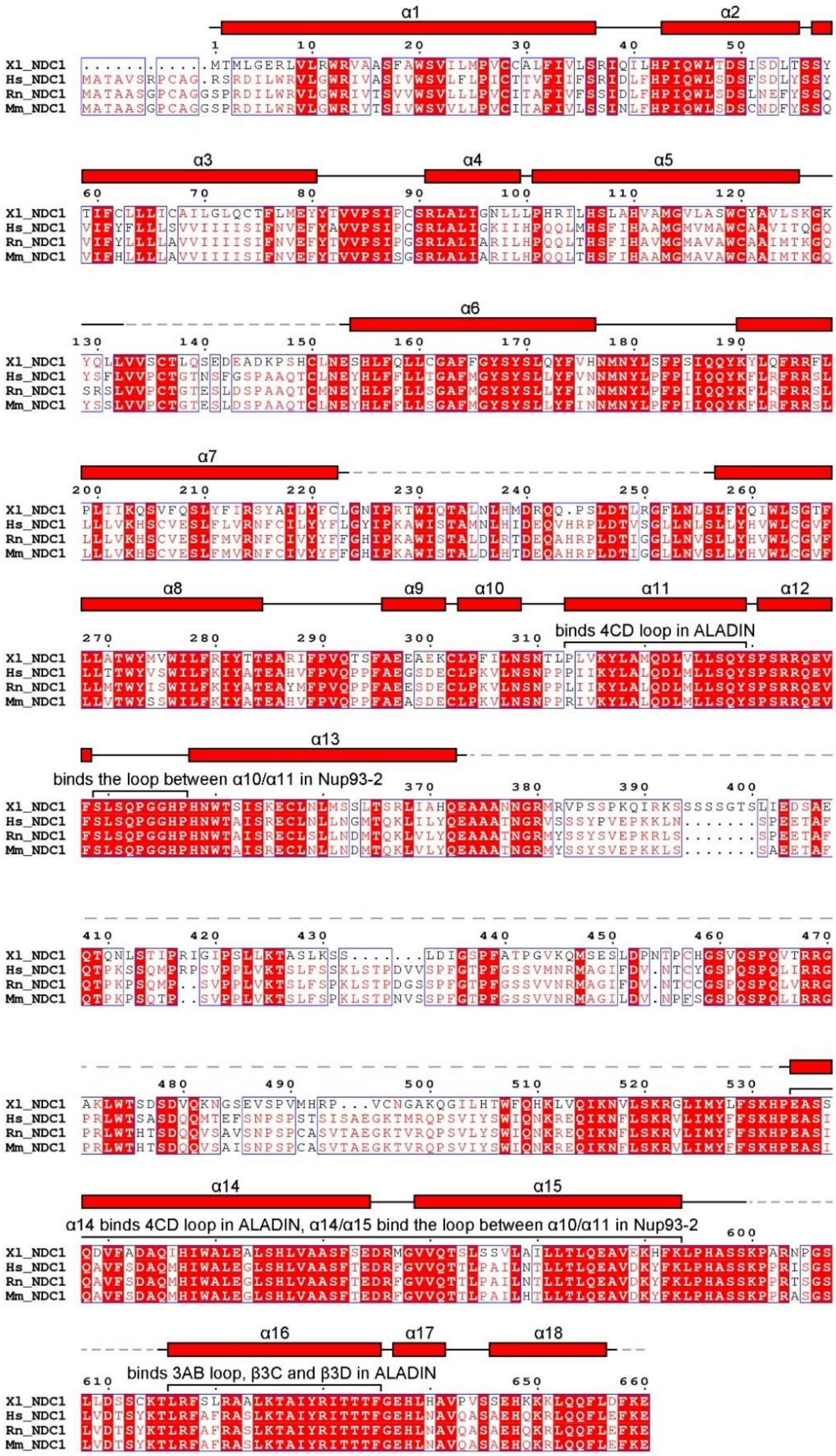


**Fig. S13 Sequence alignment among NDC1 orthologues from multiple vertebrates.** The conserved amino acids are boxed, with invariant residues highlighted in red background. The observed secondary structural elements in both NDC1 molecules are indicated above the sequences.

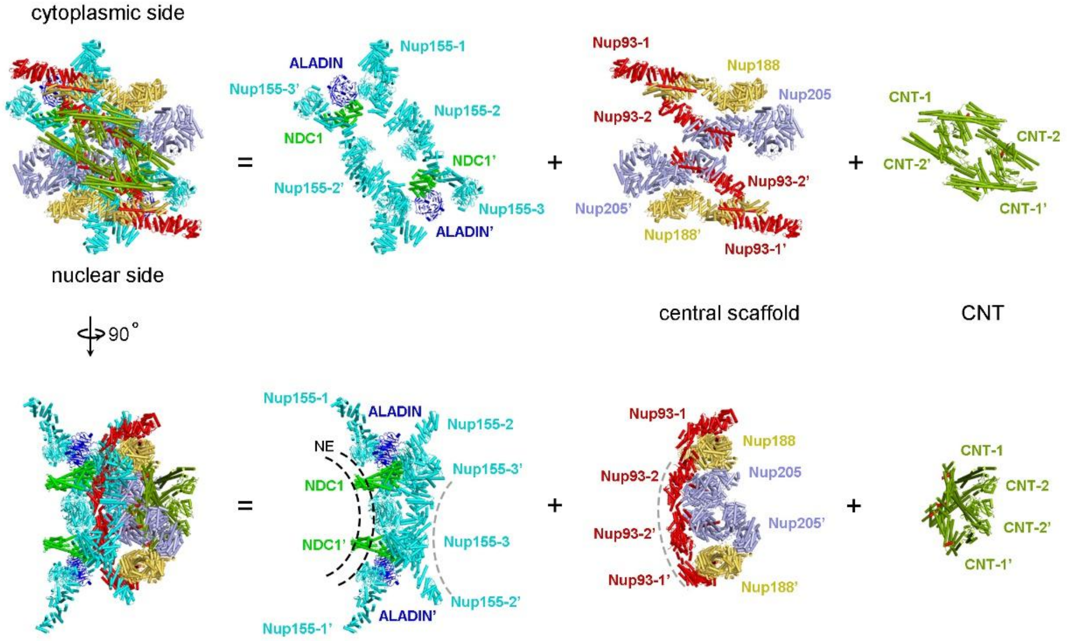


**Fig. S14 Overall structure of the IR subunit from *X. laevis* NPC.** The overall structure of the IR subunit is displayed in two perpendicular views (first column from left). For each view, the IR subunit is disseminated into three layers: 10 molecules of Nup155/ALADIN/NDC1 close to the NE (second column from left), the central scaffold of eight molecules of Nup188/Nup205/Nup93 (third column from left), and four CNTs (fourth column from left).

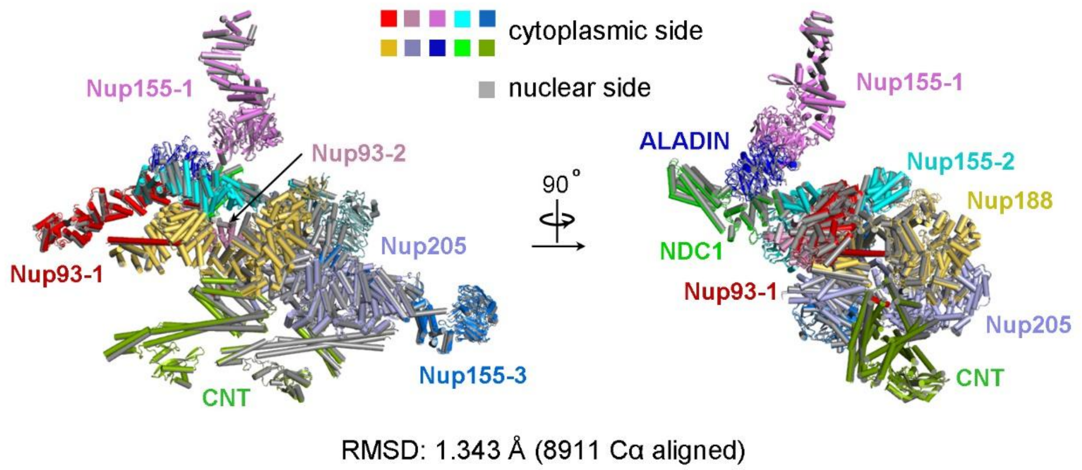


**Fig. S15 Structural superimposition of the two halves of the IR subunit.** These two halves of the IR subunit can be superimposed to each other with a root-mean-squared deviation (RMSD) of about 1.34 Å over 8,911 aligned Cα atoms. Two perpendicular views are shown.

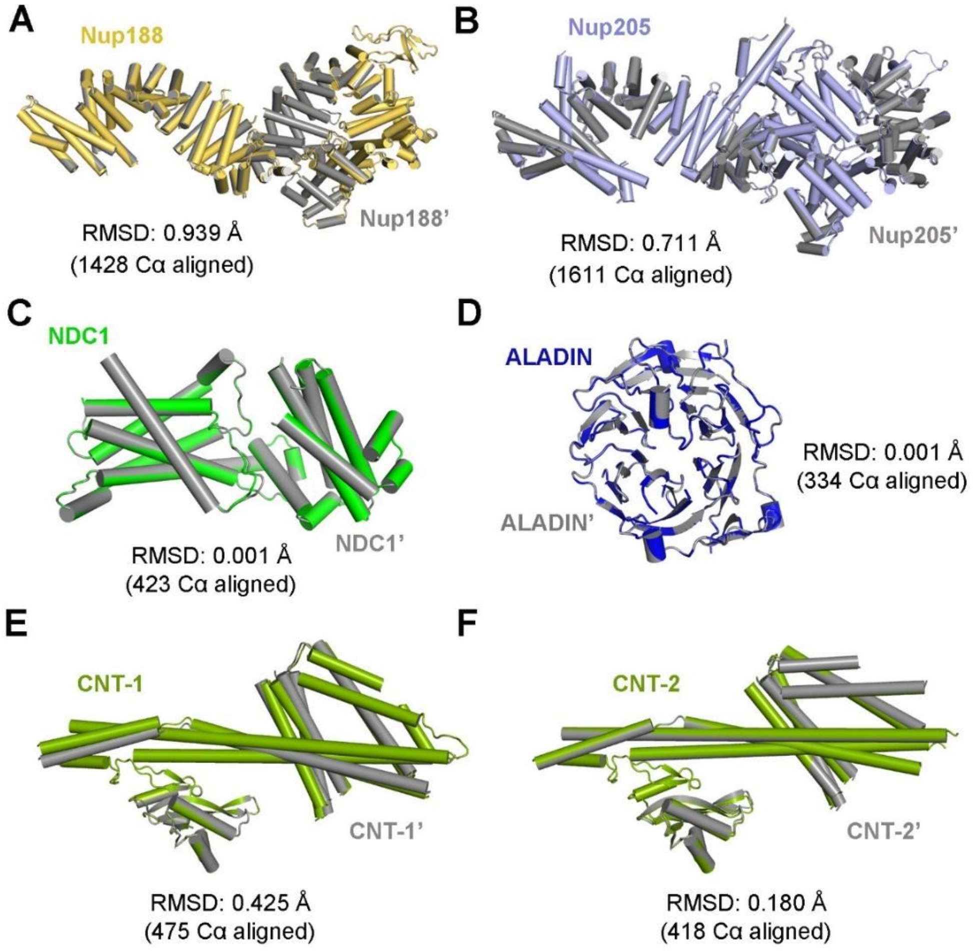


**Fig. S16 Structural comparison of select *X. laevis* nucleoporins from the IR subunit between the cytoplasmic side and the nuclear side.** (*A*) Structural comparison between Nup188 and Nup188'. (*B*) Structural comparison between Nup205 and Nup205'. (*C*) Structural comparison between NDC1 and NDC1'. (*D*) Structural comparison between ALADIN and ALADIN'. (*E*) Structural comparison between Nup93-1-associated CNT (CNT-1) and CNT-1'. (*F*) Structural comparison between Nup93-2-associated CNT (CNT-2) and CNT-2'.

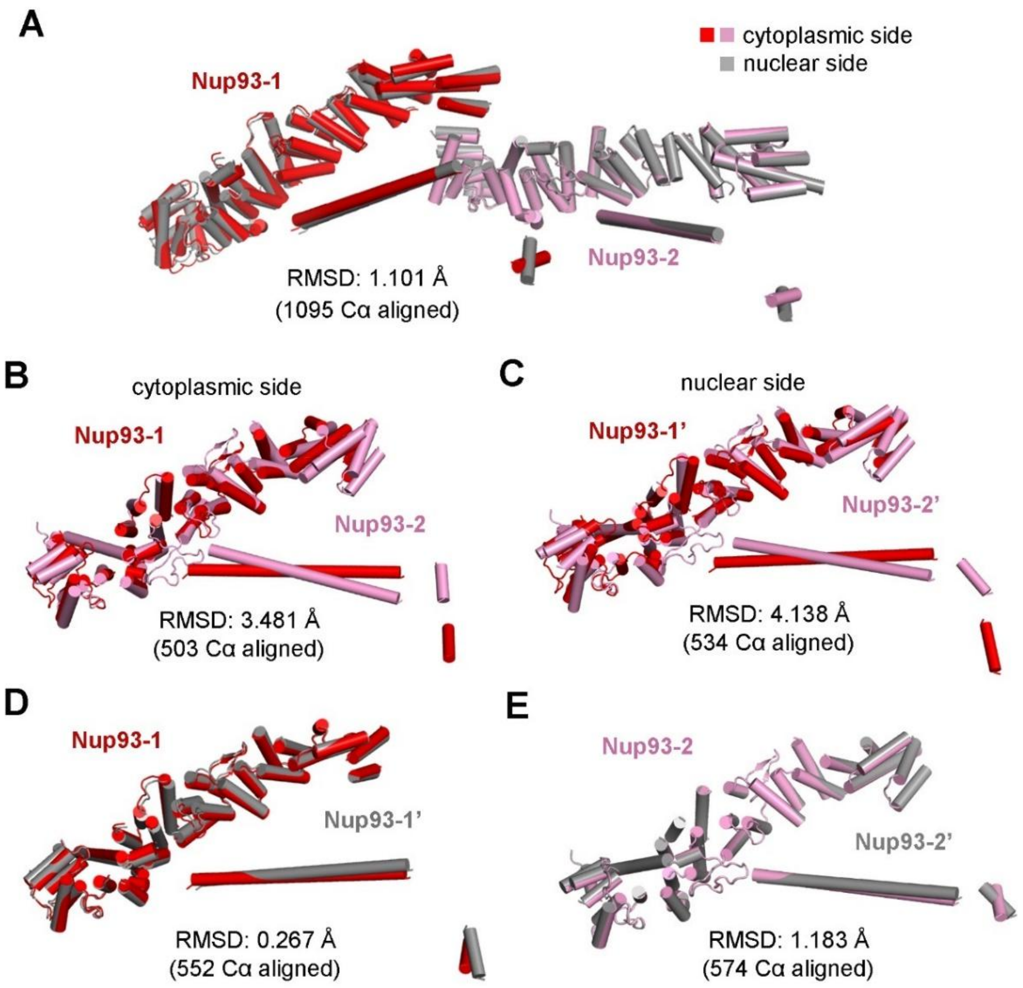


**Fig. S17 Structural comparison among Nup93 molecules within the IR subunit of *X. laevis* NPC.** (*A*) Structural comparison between the two Nup93 pairs from the cytoplasmic and nuclear sides. (*B*) Structural comparison between the two Nup93 molecules on the cytoplasmic side. (*C*) Structural comparison between the two Nup93 molecules on the nuclear side. (*D*) Structural comparison between Nup93-1 and Nup93-1'. (*E*) Structural comparison between Nup93-2 and Nup93-2'.

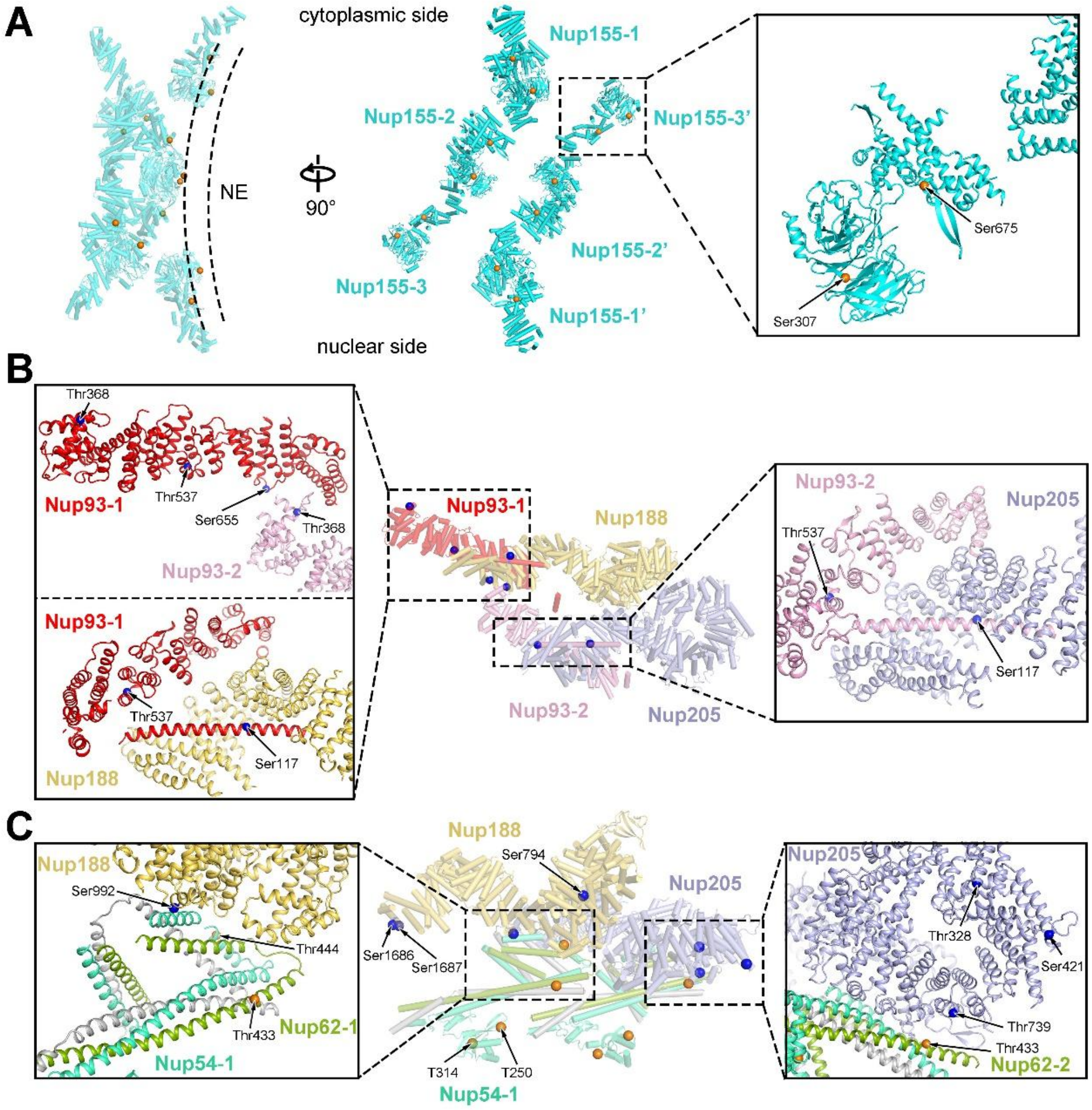


**Fig. S18 Structural mapping of phosphorylation sites on select components of the IR subunit.** (*A*) Mapping of the phosphorylation sites Ser307 and Ser675 onto Nup155. Based on our structure, both Ser307 and Ser675 face the NE. Their phosphorylation is likely to alter their interaction with the membrane, thus potentially affecting membrane localization of Nup155. (*B*) Mapping of four phosphorylation sites (Ser117, Thr368, Thr537, Ser655) onto Nup93. Phosphorylation of Ser117 may affect the interaction of the extended helix α5 with Nup205 or Nup188. Phosphorylation of Thr368 or Ser655 may alter the interface between Nup93-1 and Nup93-2. Phosphorylation of Thr537 likely changes the local conformation. (*C*) Mapping of phosphorylation sites onto Nup188, Nup205, and CNT. The phosphorylated residues include Ser794/Ser992/Ser1686/Ser1687 in Nup188, Thr328/Ser421/Thr739 in Nup205, Thr250/Thr314/Thr444 in Nup54, and Thr433 in Nup62. Their phosphorylation likely affects their mutual interaction or local conformation.

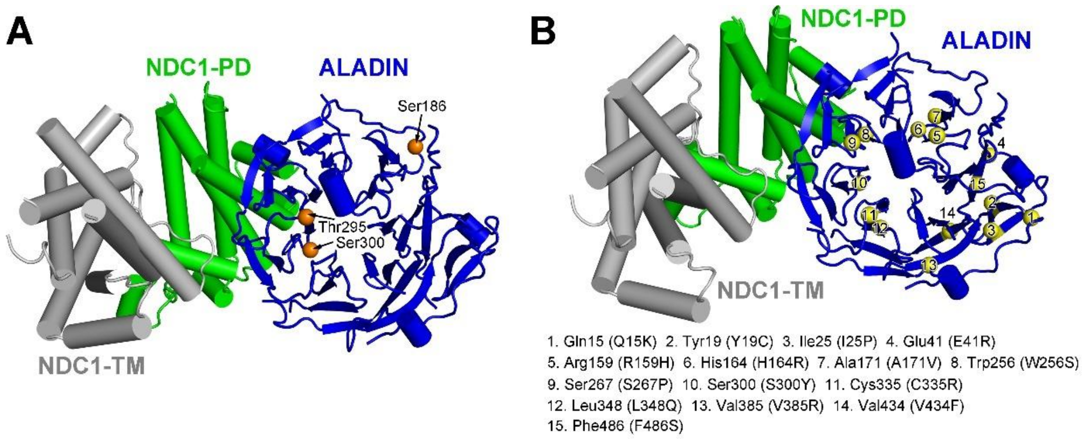


**Fig. S19 Structural mapping of phosphorylation sites and disease mutation sites on ALADIN.** (*A*) Mapping of three phosphorylation sites (Ser186, Thr295, and Ser300) on ALADIN. Phosphorylation of these sites may alter the local conformation of ALADIN. (*B*) Mapping of 15 residues in ALADIN that are mutated in the triple A syndrome (AAAS) (*18, 19*). Each of these mutations is likely to affect the local conformation and/or stability of ALADIN.

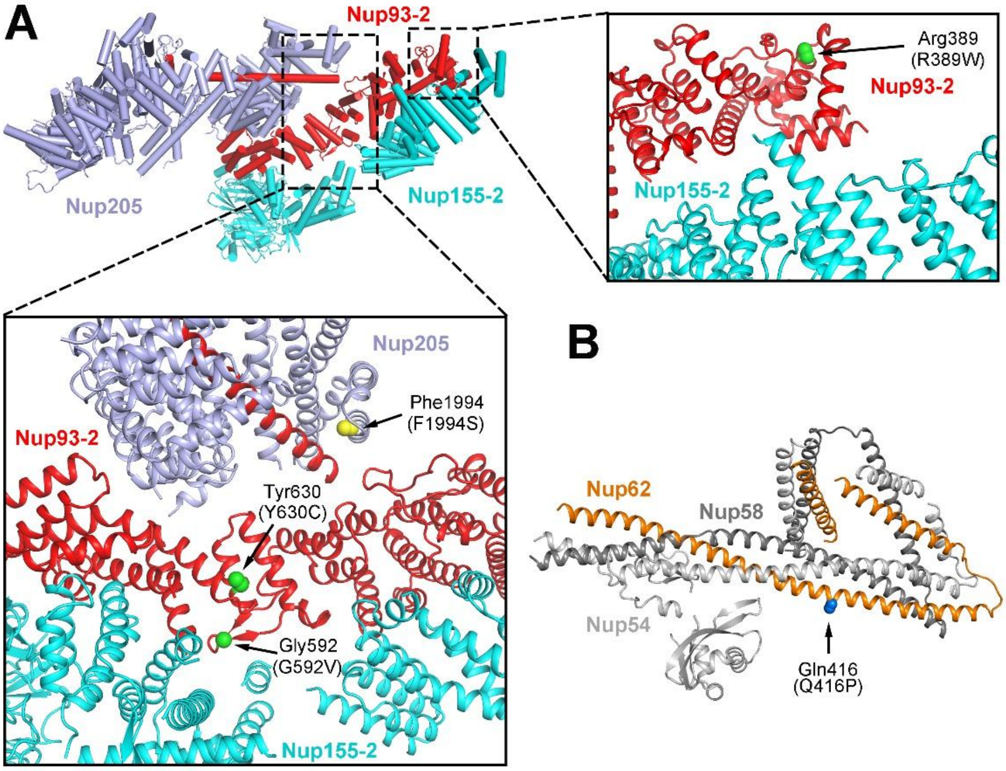


**Fig. S20 Structural mapping of disease-derived mutations onto components of the IR subunit.** (*A*) Mapping of four residues that are targeted for mutations in steroid-resistant nephrotic syndrome (SRNS) (*20*). These four residues are: Arg389/Gly592/Tyr630 in Nup93 and Phe1994 in Nup205. The mutations R389W, Y630C, G592V and F1994S may alter the local conformation. (*B*) Mapping of the disease autosomal recessive infantile bilateral striatal necrosis (IBSN) mutation Q416P in Nup62 (*21*). The mutation of Gln416 to a helix-breaking residue Pro in the middle of an extended helix may destabilize the helical conformation.

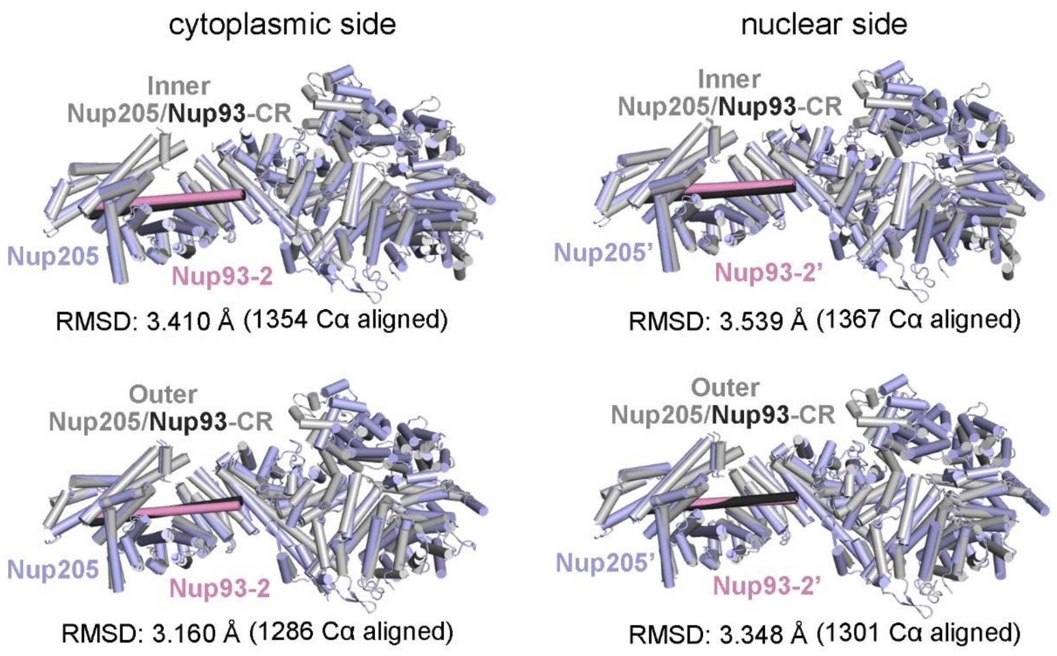


**Fig. S21 A conserved interaction between Nup93 and Nup205 in the CR and the IR.** Structural comparisons of the *X. laevis* Nup205/Nup93 pairs from the IR subunit with those from the CR subunits are shown. Comparisons of the Nup205/Nup93 pairs on the cytoplasmic and nuclear sides of the IR subunit with the Nup205/Nup93 pairs from the CR subunit are shown in the left and right panels, respectively. Comparisons between the Nup205/Nup93 pairs from the IR subunit with inner and outer Nup205/Nup93 pairs from the CR subunit are shown in the upper and lower panels, respectively.

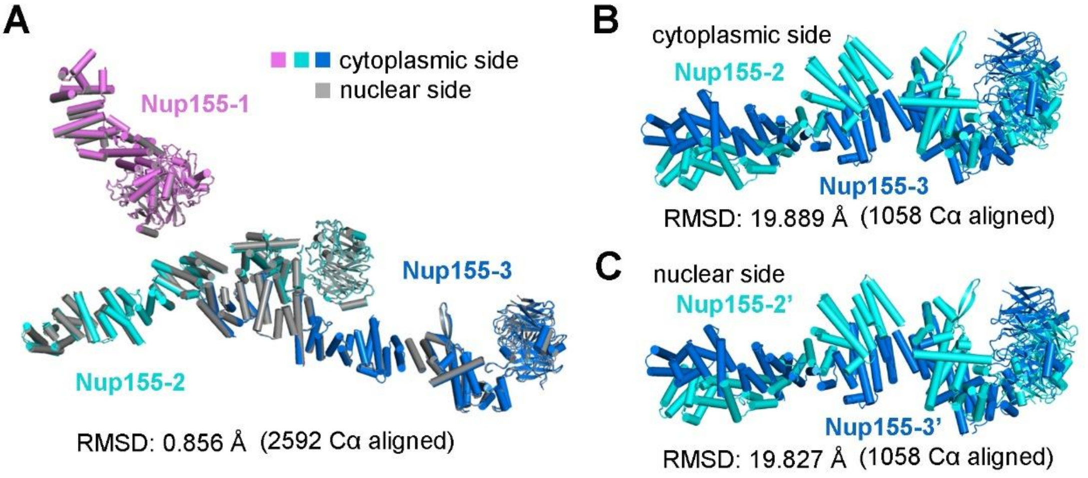


**Fig. S22 Structural comparison among six Nup155 molecules of the IR subunit.** (*A*) Structural comparison between the three Nup155 molecules on the cytoplasmic side and those on the nuclear side. The structures are nearly identical to each other, with an RMSD of 0.85 Å over 2,592 aligned Cα atoms. (*B*) Structural comparison between Nup155-2 and Nup155-3. These two structures display different local conformations, as evidenced by the RMSD of 19.9 Å over 1,058 aligned Cα atoms. (*C*) Structural comparison between Nup155-2' and Nup155-3'. Similar to the comparison between Nup155-2 and Nup155-3, these two structures also display different local conformations.

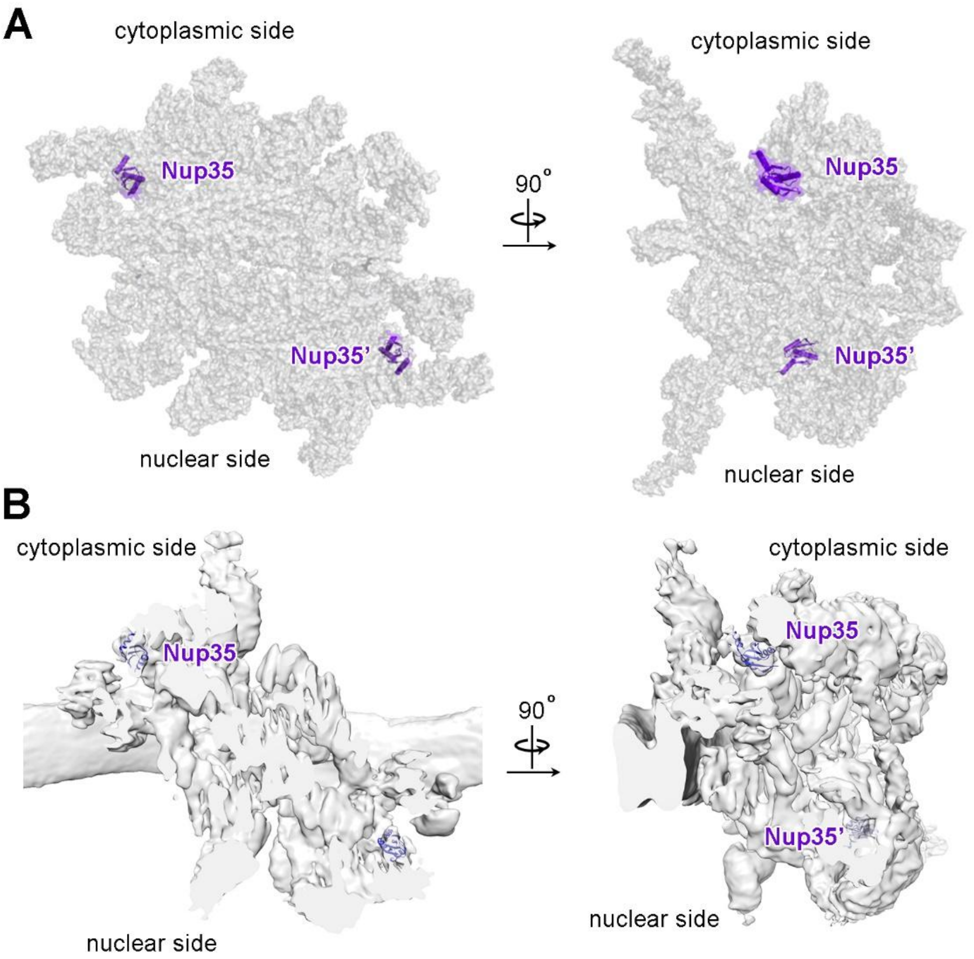


**Fig. S23  Potential locations of the RNA recognition motif (RRM) from Nup35.** (*A*) Two potential locations for the RRM domains of Nup35 are indicated in the atomic model of the IR subunit. The IR subunit is shown in transparent surface representation. (*B*) Two potential locations for the RRM domains of Nup35 are indicated in the EM map of the IR subunit.

**Table S1. Statistics of cryo-EM data collection and analysis.**

| | | | | |
|---|---|---|---|---|
| **Data collection** | | | | |
| EM equipment | Titan Krios (Thremo Fisher Scientific) | | | |
| Voltage (kV) | 300 | | | |
| Detector | Gatan K3 | | | |
| Energy filter | Gatan GIF Quantum, 20 eV slit | | | |
| Pixel size (Å) | 1.387 | | | |
| Nominal Magnification | 64,000 | | | |
| Data set | Tilt0 | Tilt30 | Tilt45 | Tilt55 |
| Exposure rate ($e^-/(s\cdot Å^2)$ ) | 19.5 | 19.5 | 19.5 | 19.5 |
| Number of frames | 32 | 37 | 46 | 56 |
| Total Electron exposure ($e^-/Å^2$) | 50 | 57 | 87 | 100 |
| Defocus range (μm) | -1.0~-4.0 | | | |
| Total Number of images | 10,040 | 7,557 | 14,107 | 14,439 |
| Selected Number of images | 8,145 | 6,171 | 10,319 | 9,112 |
| Software | AutoEMation2 | | | |
| **Reconstruction** | | | | |
| Software | RELION3.0-beta/cryoSparc | | | |
| Number of used Particles | 375,454 | 439,025 | 731,697 | 547,455 |
| Symmetry/Final resolution (Å) | C1/6.0 | | | |
| Masked regions | IR subunit core | | | |
| Number of Micrographs used | 33,747 | | | |
| Number of extracted particles | 5,223,773 | | | |
| Final number of particles | 2,093,631 | | | |
| Unmasked Resolution (Å) (0.5/0.143) | 7.5/5.5 | | | |
| Masked Resolution (Å) (0.5/0.143) | 5.6/4.3 | | | |
| Local Resolution Ranges (Å) | 50-4.0 | | | |
| Resolution Range due to anisotropy (Å) | 4.6-4.4 | | | |
| Final Resolution (Å) | 4.4 | | | |
| Map sharpening B-factor ($Å^2$) | -150 | | | |
| Accuracy of rotation (˚) | 1.61 | | | |
| Accuracy of translation (pixels) | 1.69 | | | |
| EMDB number | EMD-XXX | | | |
| **Model building** | | | | |
| Software | Coot/Chimera | | | |
| Refinement software | Phenix | | | |
| PDB code | XXX (IR subunit) | | | |
| Validation | | | | |
| R.m.s deviations | | | | |
| Bonds length (Å) | 0.002 | | | |
| Bonds Angle (˚) | 0.360 | | | |
| Ramachandran plot statistics (%) | | | | |
| Preferred | 98.00% | | | |
| Allowed | 1.92% | | | |
| Outlier | 0.08% | | | |

**Table S2. Summary of model building for the IR subunit of *X. laevis* NPC.**

| | Molecule **Vertebrates**/ Yeast | Copy No. | Length *Xenopus* | UniProt No. *Xenopus* | PDB code | Modeling | Model length | Resolution (Å) | Chain ID CR/NR side |
|---|---|---|---|---|---|---|---|---|---|
| **IR subunit** | **Nup205**/Nup192 | 2 | 2011 | Q642R6 | 7FIK | RD | ~1700 | 4.0~5.5 | A/a |
| | **Nup188**/Nup188 | 2 | 1739 | F6WXT2 | AF/4KF7/4KF8 | HM/RD | ~1500 | 4.0~5.5 | B/b |
| | **Nup93**/Nic96 | 4 | 820 | Q7ZX96 | 7FIK | RD | ~630 | 4.5~5.5 | C/E/c/e |
| | **Nup155**/Nup155 | 6 | 1388 | F6UHT0 | AF | RD | ~1150 | 4.5~6.0 | D/F/M/d/f/m |
| | **Nup62**/Nsp1 | 4 | 552 | Q6DIE3 | AF/5C3L | RD | ~170 | 5.0~6.5 | H/L/h/l |
| | **Nup58**/Nup49 | 4 | 599 | Q5EAX5 | AF/5C3L | RD | ~170 | 5.0~6.5 | I/K/i/k |
| | **Nup54**/Nup57 | 4 | 535 | K9ZTJ6 | AF/5C3L | RD | ~320 | 5.0~6.5 | G/J/g/j |
| | **NDC1**/NDC1 | 2 | 660 | Q6AX31 | AF | RD | ~430 | 5.5~6.5 | N/n |
| | **ALADIN**/- | 2 | 523 | Q6DCM0 | AF | RD | ~370 | 5.5~6.5 | O/o |

Under the column labeled "Molecule", proteins from vertebrates and yeasts are shown, respectively, with vertebrate components in bold. Under the column labeled "PDB code", AF stands for AlphaFold (model generated from AlphaFold prediction). Under the column labeled "Modeling", HM stands for homology modeling; RD stands for rigid docking and manually adjustment.

**Table S3. Summary of secondary structural elements in the structurally resolved nucleoporins of the IR subunit from *X. laevis* NPC.**

| Protein | Full length (aa) | Residues modeled | | Number of α-helices | | Number of β-strands | |
|---|---|---|---|---|---|---|---|
| | | cytoplasmic side | nuclear side | cytoplasmic side | nuclear side | cytoplasmic side | nuclear side |
| Nup205 | 2011 | 1709 | 1691 | 86 | 86 | 4 | 4 |
| Nup188 | 1739 | 1502 | 1502 | 75 | 75 | 5 | 5 |
| Nup93-1 | 820 | 635 | 635 | 32 | 32 | 2 | 2 |
| Nup93-2 | | 626 | 626 | 32 | 32 | 2 | 2 |
| Nup155-1 | 1388 | 880 | 880 | 28 | 28 | 34 | 34 |
| Nup155-2 | | 1138 | 1138 | 44 | 44 | 34 | 34 |
| Nup155-3 | | 1111 | 1111 | 44 | 44 | 34 | 34 |
| Nup62-1 | 552 | 166 | 163 | 3 | 3 | 0 | 0 |
| Nup62-2 | | 159 | 159 | 3 | 3 | 0 | 0 |
| Nup58-1 | 599 | 171 | 171 | 4 | 4 | 0 | 0 |
| Nup58-2 | | 166 | 166 | 4 | 4 | 0 | 0 |
| Nup54-1 | 535 | 318 | 314 | 10 | 10 | 6 | 6 |
| Nup54-2 | | 298 | 294 | 10 | 10 | 6 | 6 |
| NDC1 | 660 | 431 | 431 | 18 | 18 | 0 | 0 |
| ALADIN | 523 | 367 | 367 | 1 | 1 | 29 | 29 |

**References and notes:**


1. X. Zhu *et al.*, Near-atomic Structure of the Cytoplasmic Ring of the *Xenopus laevis* Nuclear Pore Complex. **Submitted**, (2021).
2. Y. Zhang *et al.*, Molecular architecture of the luminal ring of the Xenopus laevis nuclear pore complex. *Cell Res* **30**, 532-540 (2020).
3. G. Huang *et al.*, Structure of the cytoplasmic ring of the Xenopus laevis nuclear pore complex by cryo-electron microscopy single particle analysis. *Cell Res* **30**, 520-531 (2020).
4. S. Q. Zheng *et al.*, MotionCor2: anisotropic correction of beam-induced motion for improved cryo-electron microscopy. *Nat Methods* **14**, 331-332 (2017).
5. K. Zhang, Gctf: Real-time CTF determination and correction. *J Struct Biol* **193**, 1-12 (2016).
6. A. Punjani, J. L. Rubinstein, D. J. Fleet, M. A. Brubaker, cryoSPARC: algorithms for rapid unsupervised cryo-EM structure determination. *Nat Methods* **14**, 290-296 (2017).
7. S. H. Scheres, RELION: implementation of a Bayesian approach to cryo-EM structure determination. *J Struct Biol* **180**, 519-530 (2012).
8. A. von Appen *et al.*, In situ structural analysis of the human nuclear pore complex. *Nature* **526**, 140-143 (2015).
9. V. Zila *et al.*, Cone-shaped HIV-1 capsids are transported through intact nuclear pores. *Cell* **184**, 1032-1046 e1018 (2021).
10. A. P. Schuller *et al.*, The cellular environment shapes the nuclear pore complex architecture. *Nature* **598**, 667-671 (2021).
11. J. Kosinski *et al.*, Molecular architecture of the inner ring scaffold of the human nuclear pore complex. *Science* **352**, 363-365 (2016).
12. E. F. Pettersen *et al.*, UCSF Chimera--a visualization system for exploratory research and analysis. *J Comput Chem* **25**, 1605-1612 (2004).
13. T. Stuwe *et al.*, Architecture of the fungal nuclear pore inner ring complex. *Science* **350**, 56-64 (2015).
14. J. Jumper *et al.*, Highly accurate protein structure prediction with AlphaFold. *Nature* **596**, 583-589 (2021).
15. H. Chug, S. Trakhanov, B. B. Hulsmann, T. Pleiner, D. Gorlich, Crystal structure of the metazoan Nup62*Nup58*Nup54 nucleoporin complex. *Science* **350**, 106-110 (2015).
16. P. Emsley, K. Cowtan, Coot: model-building tools for molecular graphics. *Acta Crystallogr D Biol Crystallogr* **60**, 2126-2132 (2004).
17. Y. Z. Tan *et al.*, Addressing preferred specimen orientation in single-particle cryo-EM through tilting. *Nat Methods* **14**, 793-796 (2017).
18. K. Handschug *et al.*, Triple A syndrome is caused by mutations in AAAS, a new WD-repeat protein gene. *Hum Mol Genet* **10**, 283-290 (2001).
19. A. Huebner *et al.*, The triple A syndrome is due to mutations in ALADIN, a novel member of the nuclear pore complex. *Endocr Res* **30**, 891-899 (2004).

20. D. A. Braun *et al.*, Mutations in nuclear pore genes NUP93, NUP205 and XPO5 cause steroid-resistant nephrotic syndrome. *Nat Genet* **48**, 457-465 (2016).
21. L. Basel-Vanagaite *et al.*, Mutated nup62 causes autosomal recessive infantile bilateral striatal necrosis. *Ann Neurol* **60**, 214-222 (2006).
